# Shared and distinct neural signatures of feature and spatial attention

**DOI:** 10.1101/2023.08.20.554014

**Authors:** Anmin Yang, Jinhua Tian, Wenbo Wang, Liqin Zhou, Ke Zhou

**Affiliations:** Beijing Key Laboratory of Applied Experimental Psychology, National Demonstration Center for Experimental Psychology Education (Beijing Normal University), Faculty of Psychology, Beijing Normal University, Beijing, China

**Keywords:** feature attention, spatial attention, between-subject whole-brain machine learning approach, brain signature

## Abstract

The debate on whether feature attention (FA) and spatial attention (SA) share a common neural mechanism remains unresolved. Previous neuroimaging studies have identified fronto-parietal-temporal attention-related regions that exhibited consistent activation during various visual attention tasks. However, these studies have been limited by small sample sizes and methodological constraints inherent in univariate analysis. Here, we utilized a between-subject whole-brain machine learning approach with a large sample size (*N* = 235) to investigate the neural signatures of FA (FAS) and SA (SAS). Both FAS and SAS showed cross-task predictive capabilities, though inter-task prediction was weaker than intra-task prediction, suggesting both shared and distinct mechanisms. Specifically, the frontoparietal network exhibited the highest predictive performance for FA, while the visual network excelled in predicting SA, highlighting their respective prominence in the two attention processes. Moreover, both signatures demonstrated distributed representations across large-scale brain networks, as each cluster within the signatures was sufficient for predicting FA and SA, but none of them were deemed necessary for either FA or SA. Our study challenges traditional network-centric models of attention, emphasizing distributed brain functioning in attention, and provides comprehensive evidence for shared and distinct neural mechanisms underlying FA and SA.

## Introduction

The ability of visual attention to selectively focus on relevant or salient information from the environment while ignoring other stimuli is crucial for the successful functioning of various cognitive processes, including perception, memory, learning, and decision-making. Attention can be modulated across different perceptual domains, such as space (Luck et al., 1997; McAdams & Maunsell, 1999), features (Bichot et al., 2015; Liu, 2019; Maunsell & Treue, 2006), objects (Baldauf & Desimone, 2014; Chen, 2012; Roelfsema et al., 1998), time periods (Correa et al., 2006; Coull & Nobre, 1998; Denison et al., 2021), and value (Anderson, 2019; Anderson et al., 2011). These various forms of attention modulation enable individuals to allocate their cognitive resources more effectively and efficiently by prioritizing information that is most relevant to their current goals, intentions, or interests.

In spite of the importance of understanding the mechanisms of attention for improving attention and related cognitive abilities, the long-standing debate over whether different forms of visual attention have common mechanisms has remained unsolved (Chen, 2012; Maunsell & Treue, 2006; Moore & Zirnsak, 2015; Park et al., 2016; Yantis & Serences, 2003). Here we focused on two main types of visual attention: feature attention (FA) and spatial attention (SA). FA improves perception of attended features of a stimulus, while SA enhances perception at attended spatial locations.

Previous research suggested that these two forms of attentional modulation have similar neural mechanisms. Both FA and SA could strengthen the responses of the same neuron in area V4 (McAdams & Maunsell, 2000) and middle temporal area (MT) (Patzwahl & Treue, 2009). Patzwahl and Treue (Patzwahl & Treue, 2009) also found a positive correlation between the strength of feature and spatial attentional modulation at the single neuron level, implying that spatial location may not be unique, but only one of the features of a stimulus (Patzwahl & Treue, 2009). Moreover, the location and color attention-directing cues have been shown to produce largely overlapped activation in the frontoparietal network, reflecting that FA and SA shared common top-down neural control systems (Galashan & Siemann, 2017; Giesbrecht et al., 2003; Wojciulik & Kanwisher, 1999).

However, further investigations have suggested that FA and SA are distinct. These two types of attentional modulation can independently and additively affect an observer’s behavioral performance (White et al., 2015), the amplitudes of steady-state visual evoked potentials elicited in early visual areas (Anderson et al., 2011), and the responses of single V4 neurons (Hayden & Gallant, 2009). Electrophysiological recordings of neurons in area V4 revealed that SA modulated neuronal mean response gain but did not change tuning, whereas FA shifted neuronal tuning (David et al., 2008). SA often modulated spatially localized subgroups of neurons in V4, while FA could modulate neurons far apart (Cohen & Maunsell, 2011). The time course of attention also varies between FA and SA. The initial part of the visual response modulated by SA was weak, while the later part was strong, reflecting a combination of bottom-up and top-down processes. In contrast, the modulation by FA remained constant over time, reflecting a purely top-down process (Hayden & Gallant, 2005). Neurophysiological evidence from recordings in area MT implied that although spatial and feature attention may share a common top-down attention signal, they differ in their sensory normalization mechanism, as the normalization mechanism is only tuned by space, but not by feature (Ni & Maunsell, 2019). Consistent with the findings from single-neuron recordings in non-human primates, a recent study based on multivariate decoding of human MEG showed that the two forms of attention induced different representational patterns of enhancement in the occipital cortex (Goddard et al., 2022). In addition to differences in sensory modulation areas, some research also proposed that FA and SA differed in top-down control systems. A reorienting for FA resulted in a more ventral activation pattern, despite large overlap in activity between FA and SA Galashan and Siemann, 2017. Bichot and collegues (Bichot et al., 2015) identified a subregion of prefrontal cortex called the ventral prearcuate (VPA) that played a critical role in FA, because neurons in the VPA showed earlier times of feature selection than those in the frontal eye field (FEF) or inferior temporal (IT) area, and inactivation of the VPA resulted in impairment of the animal’s ability to find targets based on features while it did not affect the SA effect in FEF (Bichot et al., 2015). Further evidence supporting a distinct mechanism between the SA and FA was that SA was more sensitive to individual perceptual load than to working memory capacity, while the opposite was true for FA (Bengson & Mangun, 2018).

While previous research has explored the similarities and differences between FA and SA, the limitations of existing methods have hindered a comprehensive understanding of the neural mechanisms underlying these attentional processes. For instance, electrocorticographic (ECoG) recordings could not simultaneously take into account all the signals from the whole brain, with most studies focusing primarily on sensory areas. On the other hand, univariate analyses in functional magnetic resonance imaging (fMRI) studies have identified overlapping brain activity associated with different cognitive processes but have not provided precise representational mechanisms underlying these processes. To overcome these limitations, we employed a between-subject whole-brain machine learning-based multivariate analysis (Chang et al., 2015; Krishnan et al., 2016; F. Zhou et al., 2021) on a large sample to identify the neural representations associated with FA and SA. The approach allowed us to capture the complex and distributed patterns of neural activity across the entire brain.

Importantly, the use of machine learning algorithms enabled us to go beyond traditional univariate methods and identify fine-grained differences in the neural representations associated with different attentional processes, providing us with new insights into the functional organization of the brain regarding visual attention that were not available previously with ECoG or univariate fMRI analyses.

In this study, we employed the aforementioned machine learning approach to uncover the whole-brain multivariate patterns which could predict the conditions of FA or SA. Then we tested whether the brain signature predictive for FA was specific to FA, or it also carried information predictive for SA, and vice versa. In addition to evaluating the brain signatures at the whole-brain level, we also retrained the prediction models based on cortical networks (network level) and brain clusters (cluster level). Single-cluster and ‘virtual lesion’ analysis were applied at the cluster level to investigate the necessity and sufficiency of each cluster for FA and SA separately. Patterns of FA and SA were further compared at the voxel level to validate the distributed property of the neural underpinning.

## Materials and Methods

### Participants in the study

This study is part of an ongoing project (gene, environment, brain and behavior) (Huang et al., 2014; X. Wang et al., 2016, 2018; Y. Wang et al., 2017; Zhen et al., 2015; L. Zhou et al., 2020). 264 healthy participants (247 right-handed) were recruited from Beijing Normal University in this study. All had normal or corrected to normal vision. The study was approved by the Ethical Committee of the Beijing Normal University, and all participants were provided with written informed consent prior to the study.

Twenty-nine participants with intensive head motion (2mm in translation or 2mm in rotation from the first volume in any axis) were excluded from further analysis, leading to a final sample of 235 participants (136 female, mean age = 21.47, SD = 1.01).

### Feature attention experiment

The experiment was block-designed with 6 blocks in total (Fig 1a), three for feature task and three for conjunction task. The total block time was 32 s. The block ended with a fixation of 15 s. The order of the blocks was counterbalanced, with either feature task as the leading block and intercepted by conjunction task (F C F C F C, F: feature task, C: conjunction task) or conjunction task as the leading block (C F C F C F). Each block began with an instruction lasting 4.5 s indicating the target for the block. The sequence of letters (with size of 7.5*^◦^* * 8.5*^◦^* of visual angle) was displayed at the center of the screen against a gray background. Each trial lasted 1s with 6 letters in repaid serial visual presentation (RSVP) and with no interstimulus interval (ISI). The difference between the target and distractors could be their color, shape, or pattern. All the stimuli were either white or black. The shape of the stimuli consisted seven letters: E, H, L, O, S, T, and X, while the patterns of the stimuli came with four types: left-stripped, right-stripped, checked and wall-pattern. We designed two counterbalanced versions of RSVP. In the first version, all the distractors were black; the target in feature task was “white checked O” and the target in conjunction task was “black checked S”. In the second version, all the distractors were white; the target in feature task was “black checked O”, and the target in conjunction task was “white checked S”. This design enabled us to balance out the confounding factor of contrast while keeping both targets and distractors constant across versions. The target occurred on half of the trials with two constraints: (1) No consecutive stimuli in a RSVP were the same letter. (2) No more than three consecutive trials contained a target.

**Fig 1.**
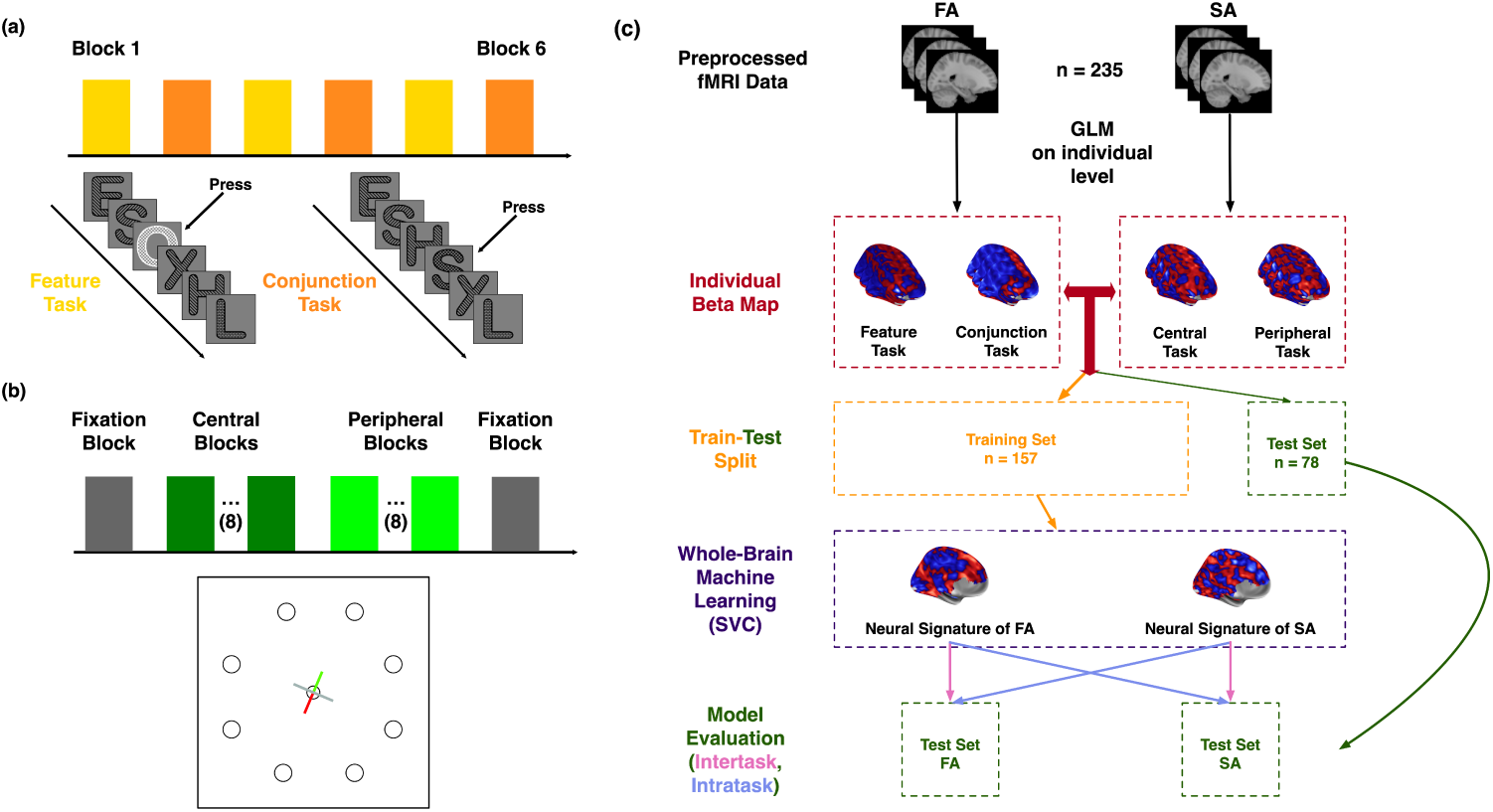
Experimental paradigms and analysis pipeline. The participants took part in both FA and SA experiments. Both experiments were block designed with two conditions. (a) In the FA experiment, serials of letters were rapidly presented. In the feature condition, participants were instructed to press the key when a white letter was presented, while ignoring other letters that were black. In the conjunction condition, participants were required to attend to both the shape and the pattern of the letter, pressing the key only when ‘black checked S’ was presented. (b) In the SA experiment, participants were instructed to press a key when the dot within the attended area decreased in size while maintaining fixation at the center of the screen. The attended area was manipulated between central and peripheral blocks. During central blocks, participants directed their attention to the central dot, whereas in peripheral blocks, their attention was focused on the dot indicated by the red arm. (c) illustrates the analysis pipeline utilized in this study. Individual beta maps (n = 235, 4 per participant) obtained from the first-level analysis were first split into training and test sets (2/3 as the training set). We employed a whole-brain multivariate machine-learning analysis using SVC to identify the distinct neural signatures of FA and SA. Model performance was evaluated using the hold-out test dataset, encompassing both intra-task and inter-task scenarios.

### Spatial attention experiment

This experiment consisted of 18 blocks (Fig 1b). The first and the last block were the fixation blocks which lasted 15 s. The remaining 16 blocks comprised spatial attention tasks: 8 blocks for central task and 8 blocks for peripheral task, depending on whether the target was the central dot or one of the peripheral dots. The 8 peripheral task blocks were further divided into 4 blocks for the left visual field condition and 4 blocks for the right visual field condition. Subjects first completed the central tasks then the peripheral tasks. The order of the left and right visual field blocks was counterbalanced. Each block began with a 3-s instruction, indicating whether the current block was a central task or a peripheral task, then followed by 20 trials. The total duration of each block was 19 s. One dot was displayed at the center of the screen and eight peripheral dots were placed evenly on a hypothetical circle whose diameter was 15*^◦^*. Against a dark gray background, the presented dots were light gray. Among them, a colored cross was located on the central dot, which had one gray arm and the other arm was half green and half red. The colored cross rotated every 4 seconds counterclockwise. In each block, the cross rotated 4 times under each visual field condition, where for the left visual field blocks, the red part of the colored cross always pointed at the dots in left visual field, and vice versa. Except for the cross, the entire display blinked on and off every 400 ms as one trial. In each display that blinked on, there was always a smaller dot on the display, either at the center or one of the four dots defined by the visual field condition. In one of the five blinks, the dot at the attended area, which was indicated by the red part of the colored cross in the peripheral task blocks or the central dot in the central task block, became smaller and reverted back to its original size on the next blink. The diameter of a normal dot was 1.3*^◦^*. The diameter of the smaller dot was 0.8*^◦^* when it was the central dot and 1.0*^◦^* when it was at the peripheral area. The task for participants was to press the key when the dot in the attended area became smaller.

### fMRI data acquisition

fMRI Scanning was performed on a 3T Siemens Trio scanner (Erlangen, Germany) at the BNU Imaging Center for Brain Research, Beijing, China. Functional images were acquired using a standard 12-channel head matrix coil, with a *T ^∗^*-weighted gradient-echo based echo-planar imaging sequence (30 contiguous interleaved 4.8mm axial slices, repetition time(TR) = 2000 ms, echo time(TE) = 30 ms, voxel size = 3.125 *×* 3.125 *×* 4.8 mm^3^, field of view (FOV) = 64 *×* 64 mm^2^, flip angle = 90*^◦^*, inter-slice gap = 0.8 mm). High-resolution structural images were acquired using a T1-weighted 3D magnetization-prepared rapid gradient-echo (3D MP-RAGE) sequence (176 slices, TR = 2530 ms, TE = 3.39 ms, inversion time = 1100 ms, voxel size = 1 *×* 1 *×* 1.33 mm^3^, FOV = 256 *×* 256 mm^2^, flip angle = 7*^◦^*. Head motion was constrained using foam pillows and extendable padded head clamps, while scanner noise was reduced by wearing earplugs.

### fMRI data preprocessing

All images were preprocessed using SPM12 (Welcome Trust Centre for Neuroimaging, http://www.fil.ion.ucl.ac.uk/spm/software/spm12/), FSL (FMRIB’s Software Library, http://www.fmrib.ox.ac.uk/fsl) and custom MATLAB functions. Preprocessing consisted of slice timing, motion correction, spatial smoothing with a Gaussian kernel (6 mm full-width at half-maximum), and high-pass temporal filtering with a 128 s cutoff. For the first-level analysis, the first four volumes of each run were discarde. First-level general linear model (GLM) analyses were conducted using SPM12. For the FA experiment, the two experimental conditions (feature condition, conjunction condition) were implemented as box-car regressors with an epoch length of 32 s, convolved with the canonical hemodynamic response function. Additionally, six motion parameters (three translation parameters and three rotation parameters) were included in the GLM as confounding variables to account for residual head motion artifacts. The GLM for the SA experiment closely mirrored that of the FA, except that the two experimental conditions (central condition, peripheral condition) were implemented as box-car regressors with an epoch length of 16 s.

### Between-subject whole-brain multivariate machine-learning analysis

We conducted a between-subject whole-brain multivariate machine-learning analysis to investigate the neural signatures of FA and SA using individuals’ beta maps derived from the first-level GLM analysis. Each individual had four beta maps, one for each attention condition (i.e., 2 conditions for FA, 2 conditions for SA). To ensure that every voxel in the prediction model possesses a numerical value, we constructed a comprehensive union mask comprising solely those voxels exhibiting beta weights across all four beta maps. This union mask was then employed to mask the beta maps, rendering them suitable for subsequent multivariate pattern analysis. For the training and testing data split, we randomly divided 1/3 of the participants (*n* = 78) as the testing dataset, using only the remaining 2/3 of the participants (*n* = 157) as the training dataset. We used support vector classification algorithm (SVC) with the linear kernel (*c* = 1) implemented by scikit-learn (Pedregosa et al., 2011) that utilizes beta maps as the training features and experiment condition as prediction targets (For FA, feature condition is denoted as 0 and conjunction condition is denoted as 1; for SA, central condition is denoted as 0 and peripheral condition is denoted as 1). Employing a linear kernel provides specific benefits, among which is its interpretability in deriving beta weights for each voxel, facilitating the prediction of the task being investigated. In addition, we also employed a PCA-Logistic algorithm (Chang et al., 2015) to validate the reliability of the results of the SVC analysis. In the PCA-Logistic method, we first used principal component analysis (PCA) to reduce the dimension of features to the number of samples (i.e., reduced to 314 features as 157 *×* 2 (number of participants *×* number of task type)). Then we used logistic regression for binary classification. The following analysis of the study was based on the beta maps from SVC analysis. To mitigate the potential bias associated with train-validation splits, we implemented a meticulous repeated 10 *×* 10-fold procedure. Within each training iteration, the complete training dataset was partitioned into 10 subsets, with 9 subsets allocated for training purposes while the remaining subset served as the validation set. This process was iterated 10 times, with the training data being re-segmented for each repetition. Notably, this meticulous procedure was repeated 10 times, ensuring robustness in our analyses. For the two attention experiments conducted, we acquired two distinct beta maps, each elucidating the predictive capacity of every voxel.

### Model Evaluation

To assess the performance of our model, we employed a diverse range of metrics, including accuracy, precision, recall, f-1 measure, and the Area under the Receiver Operating Characteristic Curve (AUC). The AUC’s scale-invariant and classification-threshold-invariant nature makes it the most suitable metric for quantifying and evaluating the effectiveness of our model. Thus, we placed particular importance on AUC as the primary evaluation measure for the model’s performance among all the metrics. Importantly, we measured the model’s performance on the hold-out test data (*n* = 78), which had been allocated prior to the training phase. Inference on model performance was performed using permutation testing via 10,000 times random shuffles of labels.

### Determining predictive voxels

In order to identify the voxels that consistently demonstrate predictive power for attentional tasks, we employed a subject-level bootstrap resampling approach. This involved generating 1,000 new training datasets by sampling with replacement. The same training procedure was then applied to each new training dataset. Consequently, for each voxel, we obtained a distribution comprising 1,000 samples. To determine the significance of voxel predictability, we then transformed this distribution into a z-score and applied a threshold to the map according to the corresponding p-value. To account for multiple comparisons, we applied a false discovery rate (FDR) correction with a threshold of *q <* 0.05. This rigorous statistical procedure enabled us to pinpoint the voxels that exhibit consistent and reliable predictive capabilities for attentional tasks.

### Univariate Analysis

To provide a benchmark for our multivariate pattern analysis, we performed univariate analysis. For each type of attentional experiment (FA and SA), we conducted two-sided matched-pair t-test on the entire sample of subjects (*n* = 235), and employed FDR correction *q <* 0.05 to control for multiple comparisons. We calculated Cohen’s *d* as an effect size measure, which served as the criteria for selecting predictive voxels (Cohen’s *d >* 0.5, denoting moderate or greater effect size).

### Network-level analysis

To discern the network-level cortical mapping distinctions between the two attentional tasks, we employed Yeo’s Atlas (Yeo et al., 2011) which leveraged resting-state functional connectivity MRI to identify seven networks comprising functionally coupled regions across the cerebral cortex. Following FDR correction, all significant voxels were classified into one of these seven distinct networks as defined by Yeo’s 1-mm liberal mask. To quantify the intensity of predictability associated with that particular network, we summed up the absolute value of all the significant voxels that fell within the network. For comparative analysis across different attentional tasks, we standardized the intensity values by expressing them as a percentage relative to the network with the highest intensity for the specific task under investigation. This approach resulted in an intensity value at 1 for one network in the respective task, with other networks as the relative intensity to the target network, which ensured consistency and enabled meaningful comparisons of intensity values across diverse attentional tasks.

To further assess the predictability of each network in both intra- and inter-task scenarios, we implemented a stepwise prediction approach that effectively controlled for the confounding influence stemming from voxel count disparities among networks.

Starting with 10 voxels, we progressively increased the number of voxels included in the prediction model until all voxels within the network of interest were incorporated. At each step, a fixed number of voxels within the network were randomly selected as features for training SVCs to predict the conditions of either FA or SA. The performance of the models was then evaluated on the hold-out test dataset for both intra-task and inter-task scenarios. The training and testing protocols followed the same methodology as employed in the whole-brain analysis. To ensure robustness of our findings, we repeated the training and testing procedures 100 times at each step.

### Cluster-level Analysis

We used the watershed image segmentation algorithm (Fedorenko et al., 2010; Meyer, 1994; Zhen et al., 2015) to partition the significant voxels, after FDR correction, into separate clusters within the beta maps. By isolating these clusters, we were able to delve into their individual contributions to attentional processes. Specifically, we utilized single-cluster prediction analysis and ‘virtual lesion’ analysis to examine the predictive sufficiency and necessity of individual clusters in attentional tasks, respectively. In the single-cluster prediction analysis, we focused on isolating the cluster of interest and applied the same model training procedures employed in the whole-brain analysis. Subsequently, we assessed the model’s performance in both intra- and inter-task scenarios. If the cluster of interest is indeed sufficient in predicting attentional tasks, we would anticipate an AUC score exceeding chance level. This analysis provides valuable insights into the independent predictive contribution of the isolated cluster in attentional processing. In the ‘virtual lesion’ study, we performed an analysis wherein we excluded the cluster of interest from the whole-brain signature and proceeded to retrain the classifier. If the performance of the newly trained prediction model drops to chance level, it provides compelling evidence that the lesioned cluster plays a necessary role in the specific attentional task. Again, we implemented a gradual single-cluster prediction and ‘virtual lesion’ analysis with a step size of 10 voxels, to exclude the potential confounding effect of voxel number variations across clusters when comparing their influence on the change in model predictability. In each step, we randomly selected the same number of voxels from each cluster, and this process was repeated 1,000 times to retrain the prediction models. This procedure allowed us to ensure comparable model complexity across different clusters, effectively controlling for any voxel number variations. The stepwise study also incorporated the analysis of whole-brain signatures.

### Voxel-level Analysis

To investigate the joint normalized distribution of the neural signatures of FA and SA, we utilized a visualization technique to represent them in a 2-D space. To ensure comparability among voxels within neural signatures, we first normalized the weights within each signature independently. Thus, for each voxel, the x-axis represented its normalized weight for FA, and the y-axis represented its normalized weight for SA. Next, we divided the 2-D space into octants, with each two octants corresponding to a distinct conceptual category: FA dominant, SA dominant, same pattern, and opposite pattern. To incorporate the cortical relationship among these octants, we calculated the coverage ratio of significant voxels falling into their respective networks. These ratios were then normalized by expressing them as proportions relative to the highest coverage ratio observed within the network. This normalization procedure allowed us to account for variations in the coverage across different networks and facilitated fair comparisons among them. By employing this approach, we gained insights into the distribution patterns and cortical relationships among the various categories within the joint FA and SA signatures.

## Results

### Participant’s behavioral performance in FA and SA tasks

235 participants underwent both FA and SA tasks with fMRI scanning (Fig 1). In the FA task, participants were shown with series of letters, where they need to respond to a white letter for the feature condition and a combination of shape and color for the conjunction condition. In the SA task, participants should orient their attention either to the center or the pheriphral region to respond when the size of the dot changed.

Fig 2a showed the sensitivity (*d^′^*) and response time (RT) in the FA and SA experiments (details see Supplementary Table 1). In the FA experiment, both sensitivity and RT were significantly different between feature condition and conjunction condition (two-sided paired sample t-test: *d^′^*: *t*(234) = 55.18, *p <* 0.001, Cohen’s *d* = 4.92; RT: *t*(234) = *−*36.29, *p <* 0.001, Cohen’s *d* = 2.75), supporting conjunction condition being more difficult. Likewise, in the SA experiment, the peripheral condition was more challenging than the central condition, yet with weaker effect size (two-sided paired sample t-test: *d^′^*: *t*(234) = 16.74, *p <* 0.001, Cohen’s *d* = 1.33; RT: *t*(234) = *−*2.78, *p* = 0.008, Cohen’s *d* = 0.17). The less sensitivity and slower RT in difficult conditions was consistent with the behavioral performance discovered in Wojciulik and Kanwisher (1999).

**Fig 2.**
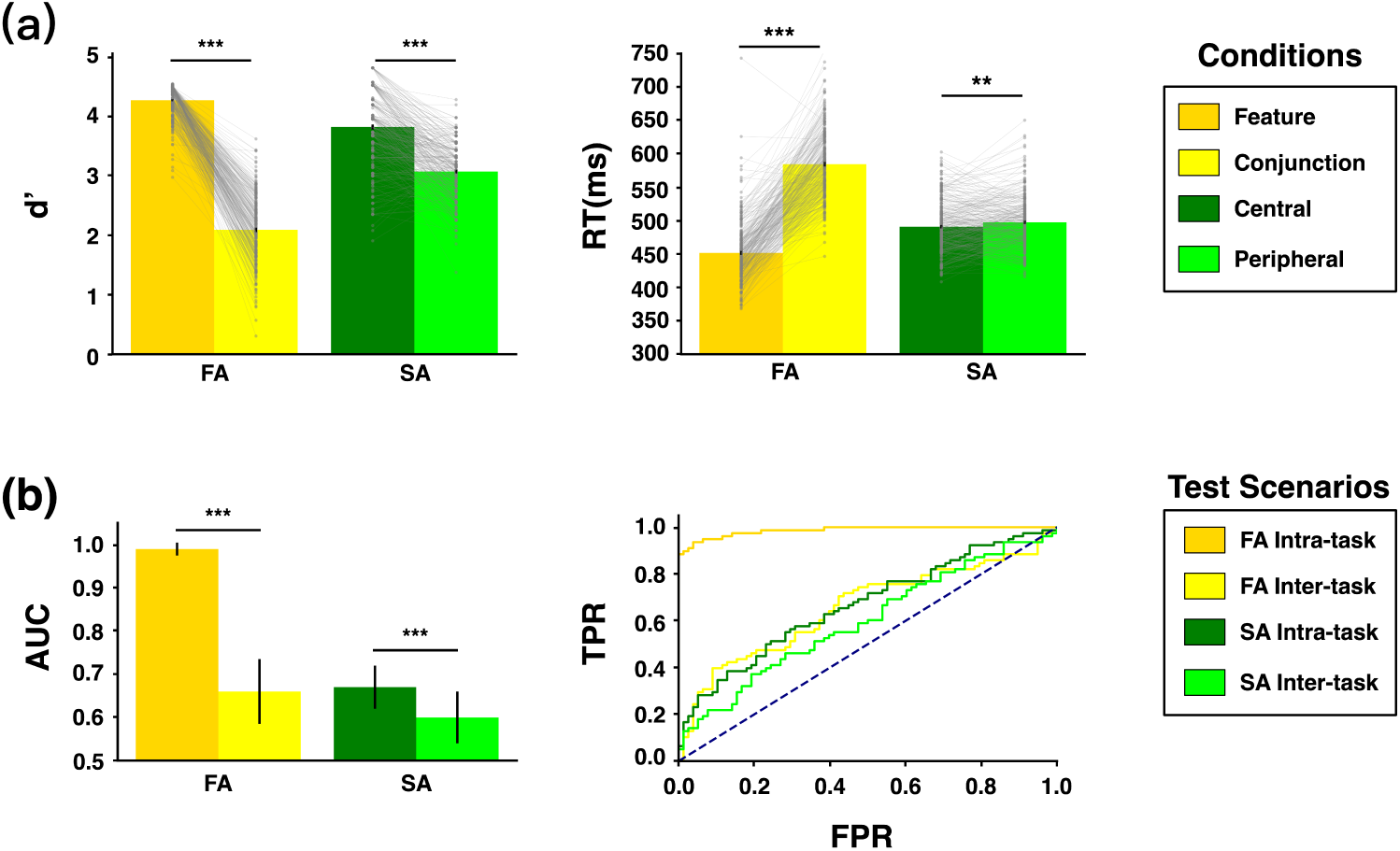
Behavioral results and model performance. (a) presents the behavioral results of the FA and SA experiments. The *d^′^* index quantifies participants’ discriminability of signals from distractors, while Reaction Time (RT) measures the time elapsed from signal appearance to the subject’s key press response, serving as metrics of task difficulty. The error bar represents the standard error (SE) of measurement. (b) Model performance of the neural signatures of FA and SA was assessed using the AUC metric, with error bars representing the 95% confidence interval (CI) obtained through 10,000 bootstrap iterations. The receiver operating characteristic curve (ROC) is depicted on the right, showing true positive rate (TPR) and false positive rate (FPR). Asterisks indicate the p-value obtained from matched-sample two-tailed t-test: ***, *p <* 0.001; **, *p <* 0.01.

### The FAS and SAS were capable of mutually predicting each other’s behaviors

We applied the Support Vector Classifier (SVC) with a linear kernel to unveil the brain signatures of FA (FAS) and SA (SAS) separately. Each attention type’s brain signature consisted of the voxel weights that reflected their respective contribution to predicting the attention condition. To evaluate the robustness and mutual predictability of these brain signatures, we conducted tests on a hold-out dataset (*n* = 78) including both the intra-task scenario (e.g., using the FAS to predict FA conditions) and inter-task scenario (e.g., using the FAS to predict SA conditions). The prediction performance was quantified using the Area Under the Curve (AUC) metric, as illustrated in Fig 2b (for other model performance metrics see in Supplementary Table 2). First, both brain signatures exhibited prediction performance above chance in intra-task scenario (FA intra-task: AUC = 0.99, bootstrapped 95%CI = [0.97, 1.00], permutation test one-tailed *p <* 0.001; SA intra-task: AUC = 0.67, bootstrapped 95%CI = [0.62, 0.72], permutation test one-tailed *p <* 0.001). We also observed that the SAS exhibited comparatively lower prediction performance within intra-task scenario when compared to the FAS. One possible explanation for this discrepancy could be the larger variation in task difficulty between conditions in the FA experiment. To test whether the difference of task difficulty between FA and SA conditions could affect the result of the FAS and SAS, we selected 84 subjects as a subset of the total training set, whose differences of *d^′^* in two conditions were similar between FA and SA (two-sided paired sample t-test: *t*(83) = 1.824, *p* = 0.070). We employed the same procedure to acquire the FAS and SAS and found no major impact of task difficulty on the FAS and SAS compared to the full training set(dice coefficients were 0.62 and 0.91 for FA and SA respectively).

More interestingly, both FAS and SAS exhibited prediction performance above chance level when applied to inter-task scenarios (FA inter-task: AUC = 0.66, bootstrapped 95%CI = [0.58, 0.73], permutation test one-tailed *p <* 0.001; SA inter-task: AUC = 0.60, bootstrapped 95%CI = [0.54, 0.66], permutation test one-tailed *p <* 0.001). These findings suggested the presence of shared neural components within the FAS and SAS.

However, it was also observed that the SAS outperformed the FAS when tested on the SA dataset (permutation test one-tailed *p* = 0.012), indicating a higher sensitivity of the SAS in predicting SA conditions compared to the FAS. Conversely, the FAS demonstrated better predictive capabilities on the FA dataset than the SAS (permutation test one-tailed *p <* 0.001). In addition, both FAS and SAS exhibited a significant drop in performance from the intra-task scenario to the inter-task scenario, suggesting the existence of distinct components between the FAS and SAS (permutation test one-tailed, FAS: *p <* 0.001, SAS: *p* = 0.038).

To verify the robustness of the signatures, we additionally employed a PCA-Logistic algorithm and compared the signatures identified by both approaches (for details see Methods). Remarkably, the signatures derived from both approaches exhibited a high degree of concordance, as evidenced by a significant correlation in weight (For FA, r=0.76, 95%CI = [0.75, 0.76], *p <* 0.001; For SA, *r* = 1.00, 95%CI = [1.00, 1.00], *p <* 0.001, see Supplementary Figure 1). These results provided compelling evidence for the robustness of the identified signatures.

### FAS and SAS exhibited both shared and distinct components

Using a linear kernel in the SVC models, we identified the FAS and SAS (Fig 3a). To ensure reliable voxel predictability, we utilized bootstrap resampling and employed FDR correction with a threshold of *q <* 0.05 for multi-comparison correction. The FAS and SAS manifested both shared and distinct components. As illustrated in Fig 3a, the brain regions within these signatures could be categorized into three categories: those specific to FA, those specific to SA, and the conjunction regions. The conjunction regions were widely distributed across frontoparietal areas and visual areas, including bilateral IPTO (junction of intraparietal and transverse occipital sulci), left aSMG (anterior supramarginal gyrus), right aCG (anterior cingulate gyrus), bilateral PcC (precuneus cortex), right OFG (occipital fusiform gyrus), right PrG (precentral gyrus) and right OcP (occipital pole), etc. Regarding the regions specific to FA, we observed activations in the right AG (angular gyrus), bilateral SFG (superior frontal lobe), bilateral COpC (central opercular cortex), and left FOpC (frontal opercular cortex), etc. In contrast, we did not find any regions specific to SA.

**Fig 3.**
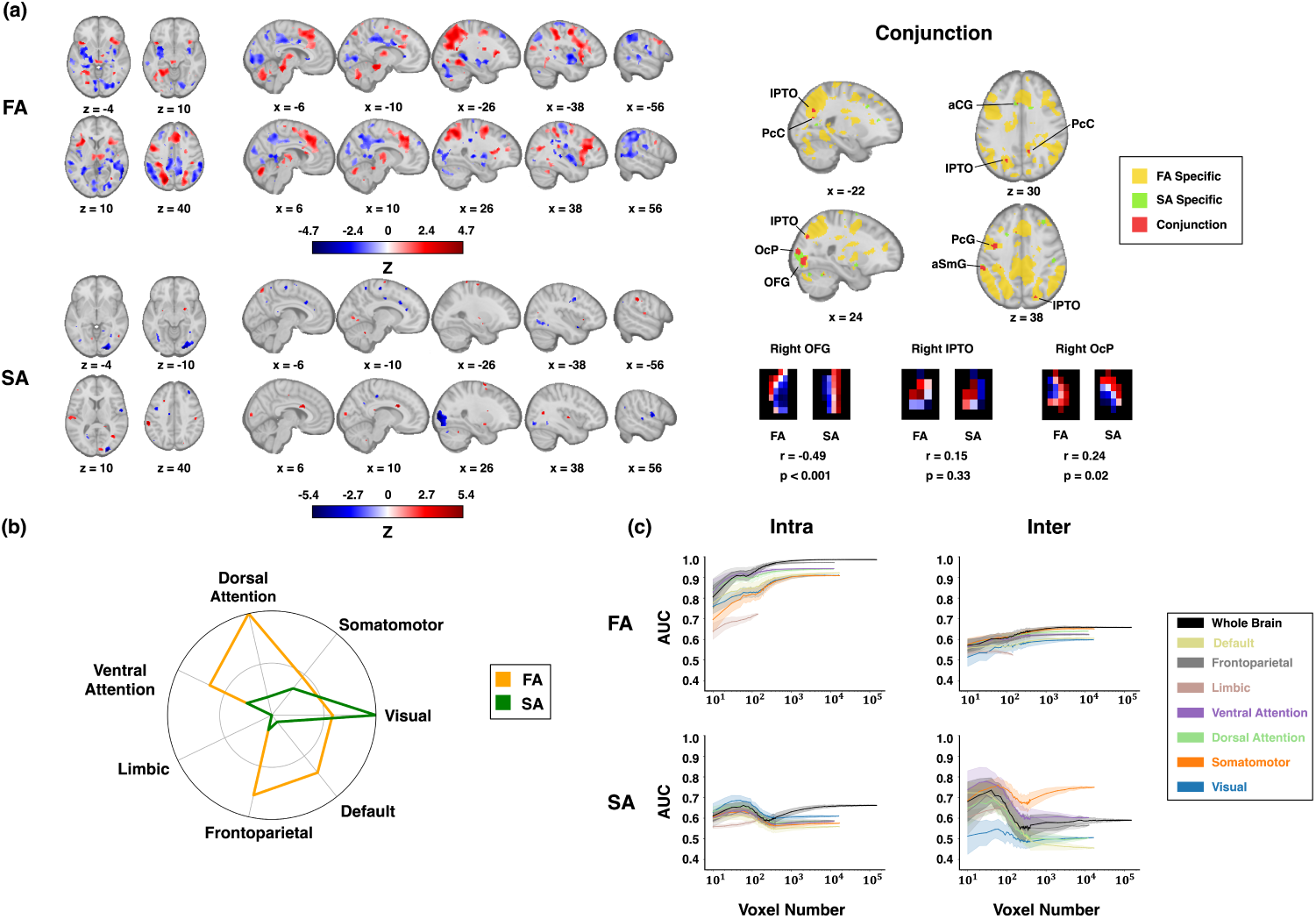
Neural signatures of FA and SA. (a) depicts the brain signatures of FA and SA with predictive voxels determined by bootstrap over 1,000 times and corrected by FDR at *q <* 0.05. Conjunction plots illustrate the voxels falling into three categories: FA Specific, SA Specific, and the overlapping regions between FA and SA. IPTO: intraparietal/transverse occipital sulci, PcC: Precuneus Cortex, aCG: anterior Cingulate Gyrus, OcP: Occipital Pole, OFG: Occipital Fusiform Gyrus, aSmG: anterior Supramarginal Gyrus. (b) The intensity of predictive voxels in FAS and SAS is presented across 7 cortical resting-state networks. (c) The predictivity of each network across the two tasks is depicted, with an equal number of voxels in each step.

In addition, we conducted a univariate analysis similar to a previous study by Wojciulik abd Kanwisher (Wojciulik & Kanwisher, 1999), who identified overlapping regions such as IPTO and aIPS (anterior intraparietal sulcus). Consistent with their findings, we also found overlapping regions at IPTO, FEF (frontal eye field) and SPL (superior parietal lobe) (see Supplementary Figure 2). These results provided robust validation of our experimental design and data.

However, it is crucial to acknowledge that while univariate analysis can reveal brain regions that co-activate during FA and SA, assuming that the whole overlapping region represents a homogeneous common neural substrate underlying different attentional tasks might oversimplify the complexity of the underlying neural mechanisms. By employing multivariate machine-learning analysis, we have uncovered that even within the brain regions identified through conjunction analysis in the univariate approach, distinct patterns could exist. This highlighted the importance of adopting a multivariate approach to capture the nuanced variations and heterogeneity within these shared brain regions. Specifically, we found that the right OFG (*r* = *−*0.49, *p <* 0.001) and left PrG (*r* = *−*0.39, *p <* 0.001) exhibited the opposite patterns, whereases the right OcP showed a similar pattern (*r* = 0.24, *p* = 0.02)(Fig 3a). On the other hand, other conjunction regions, such as bilateral IPTO (left: *r* = *−*0.11, *p* = 0.73; right: *r* = 0.23, *p* = 0.33), did not exhibit correlated patterns (also refer to Supplementary Table 3).

To assess the distribution of the two brain signatures across established intrinsic cortical networks, we computed voxel weight sums for each network using Yeo’s atlas (Yeo et al., 2011). As depicted in Fig 3b, the magnitude of a specific brain network was determined by the proportion of weight sums from significant voxels within that network, relative to the network with the highest intensity. This analysis revealed distinct involvement patterns of the FAS and SAS in brain networks defined by Yeo’s atlas. This quantification enabled us to assess the relative contribution of each network in terms of the weight distribution of the significant voxels. The most prominent regions associated with the FAS were the dorsal and ventral attention networks, whereas the SAS was predominantly distributed within the visual network.

In addition, conducting the prediction analysis with stepwise controlled voxel number further confirmed the dissociation among cortical networks (Fig 3c). As the number of significant voxels in SA was considerably smaller than in FA, increasing the voxel count in the prediction models could lead to the incorporation of additional noise. This phenomenon likely explained the observed decline in performance within each network, which typically occurred at around 100 voxels.

### Single-Cluster and ‘Virtual Lesion’ analyses revealed similar and distinct representation patterns in FA and SA, along with their signatures’ distributed nature

The multivariate machine-learning analysis and the univariate analysis have revealed the distributed activations throughout the brain elicited by both FAS and SAS. Then how sufficient and necessary each cluster of the signatures was to predict FA and SA? To address the question, we conducted single-cluster analysis and ‘virtual lesion’ analysis. Using the watershed algorithm with the minimum size at 10 voxels, we first defined the clusters within the FAS and SAS. The conjunction clusters were primarily found in fronto/parietal, temporal/occipital, and limbic areas. Their prediction performance in both single-cluster and ‘virtual lesion’ analysis was presented in Table 1 and Table 2. Supplementary Table 4 showed the performance of all the clusters in FAS.

**Table 1.**
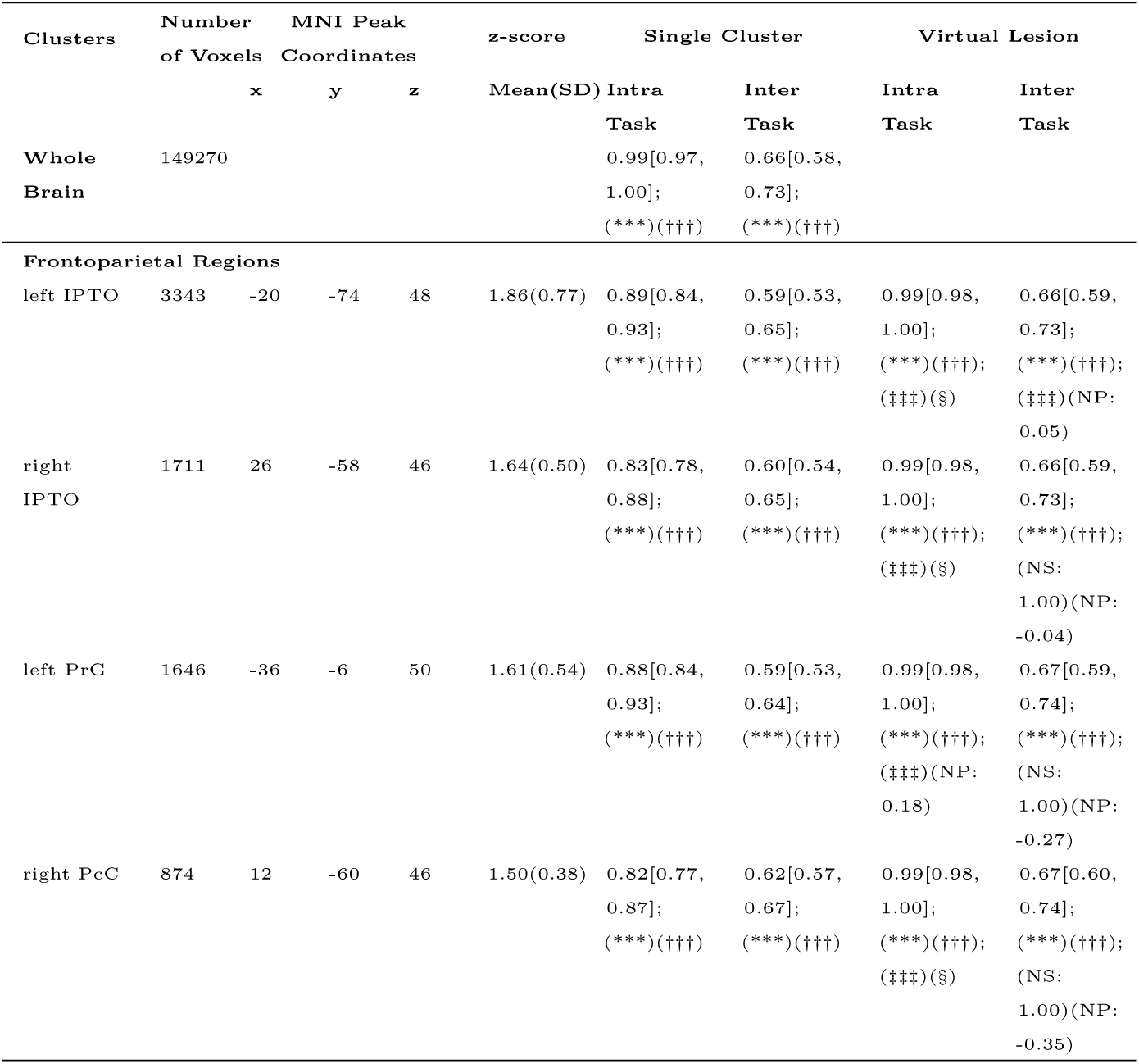
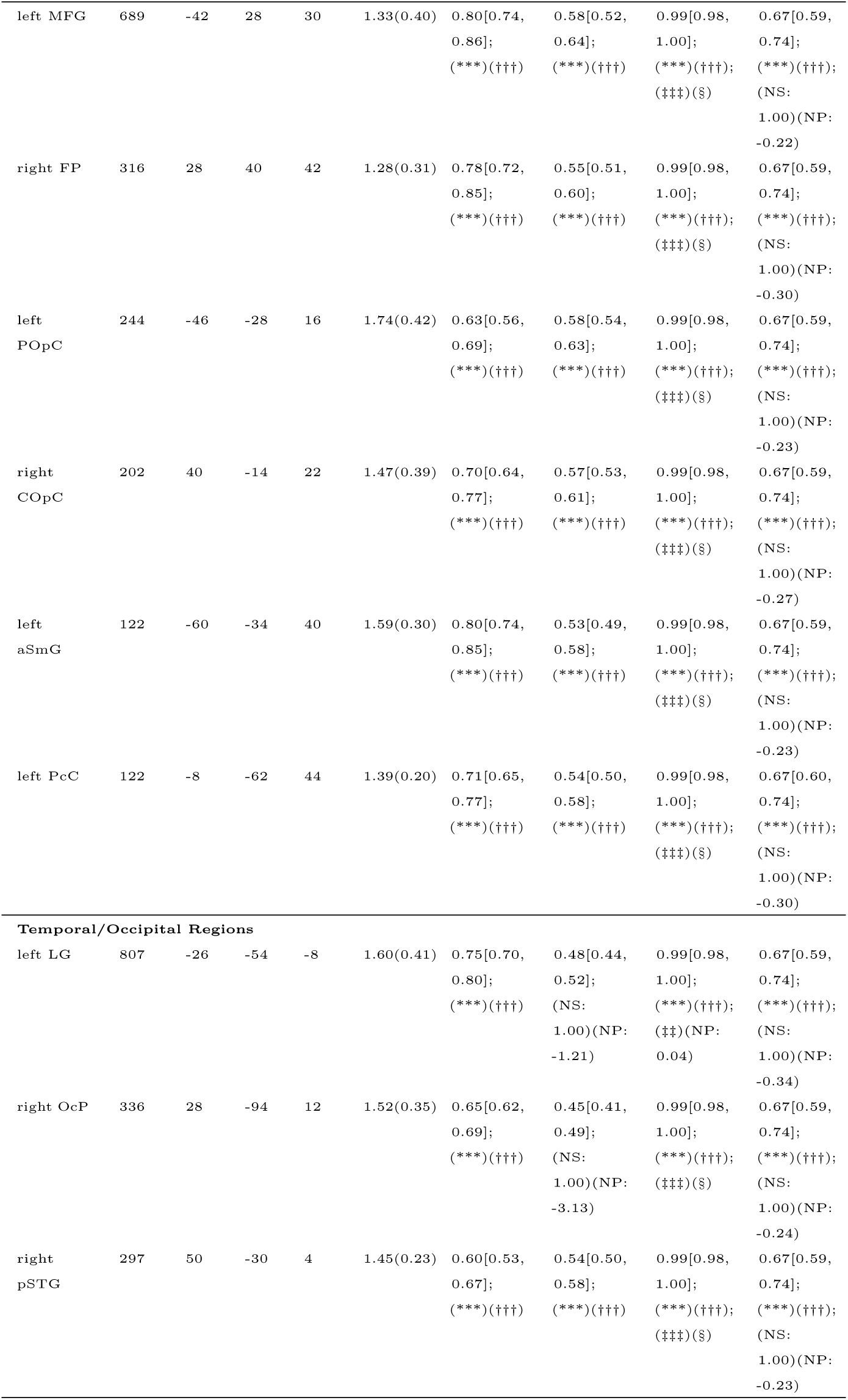
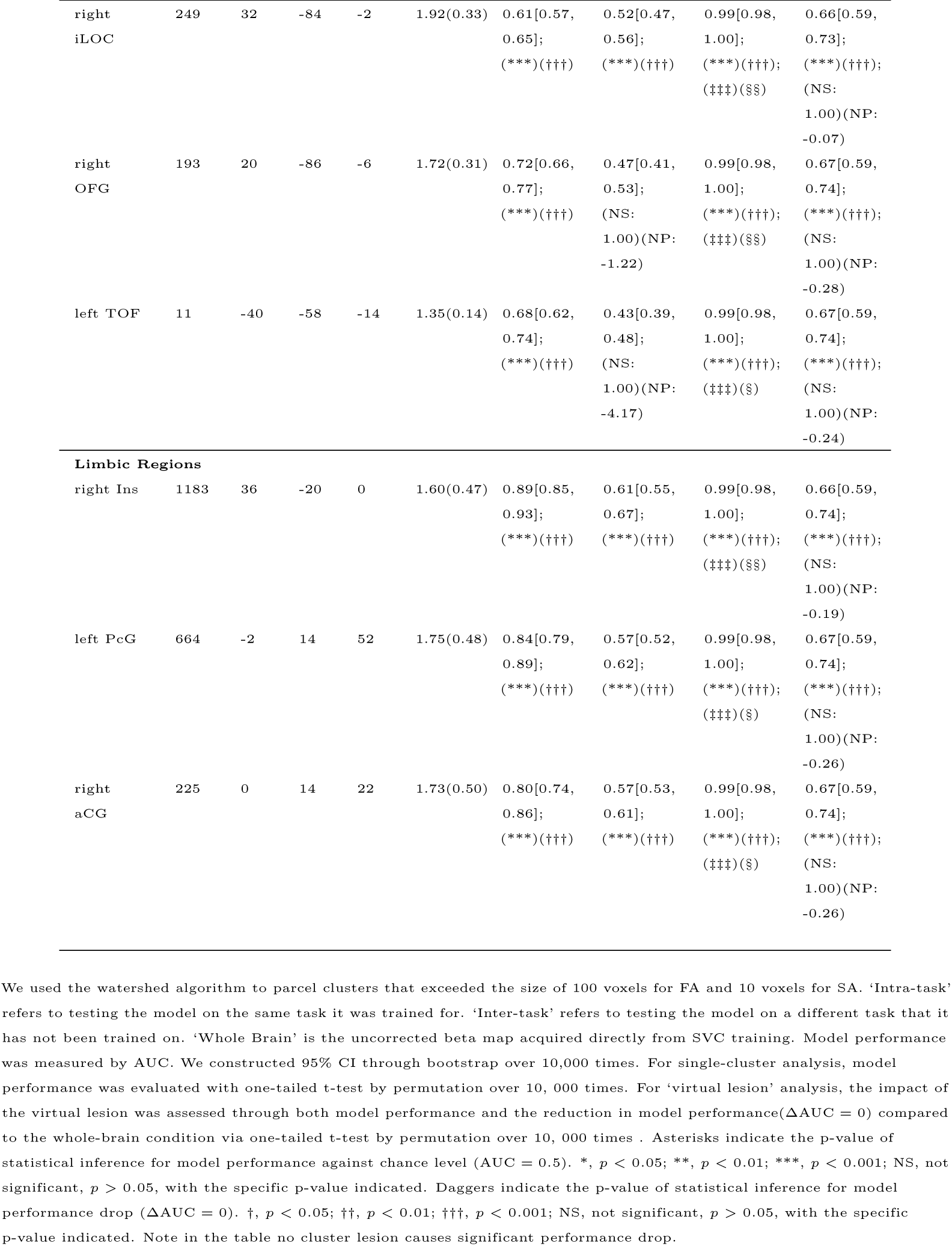
Single-cluster and ‘Virtual Lesion’ analysis of FA.

**Table 2.**
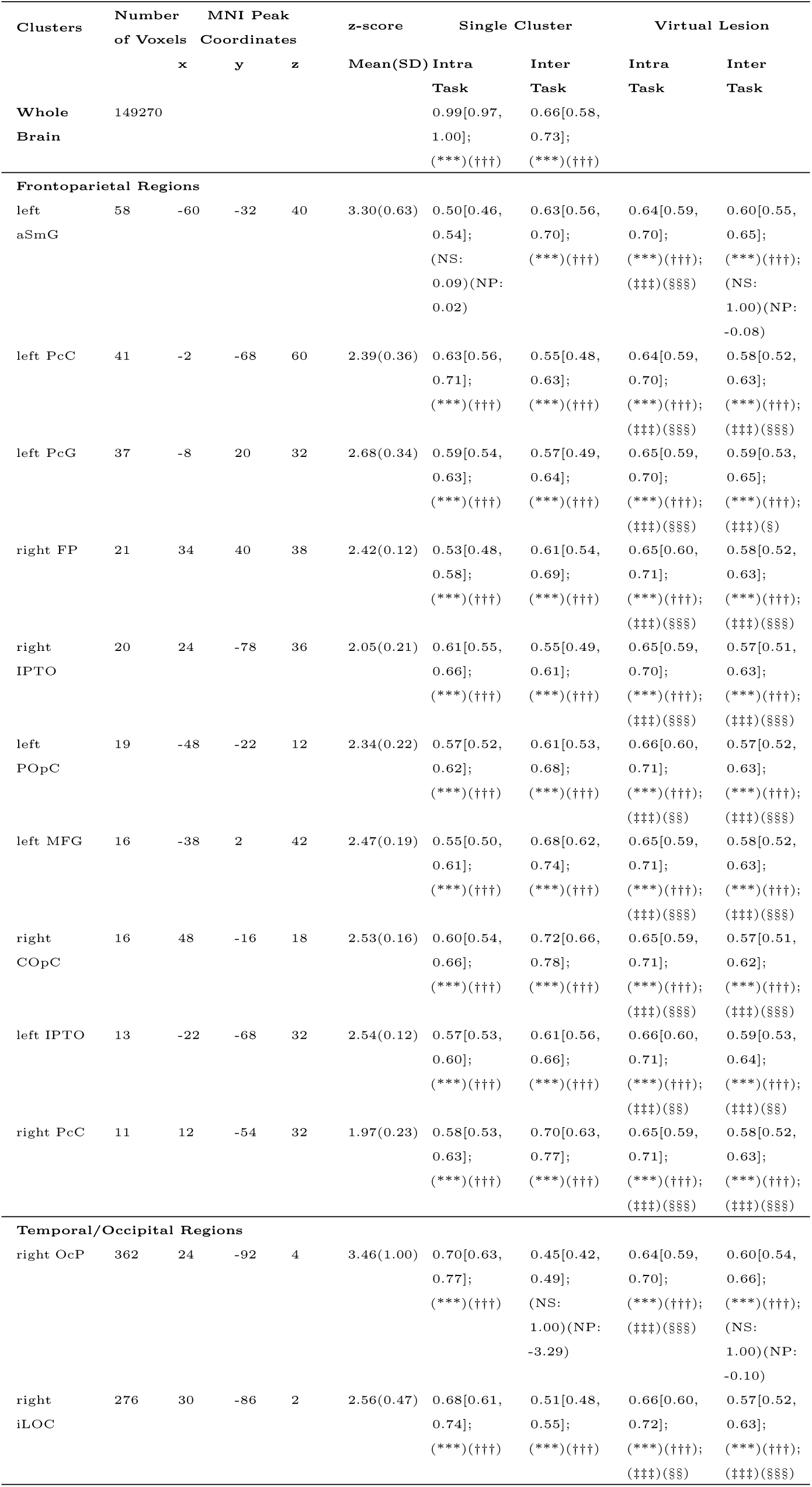
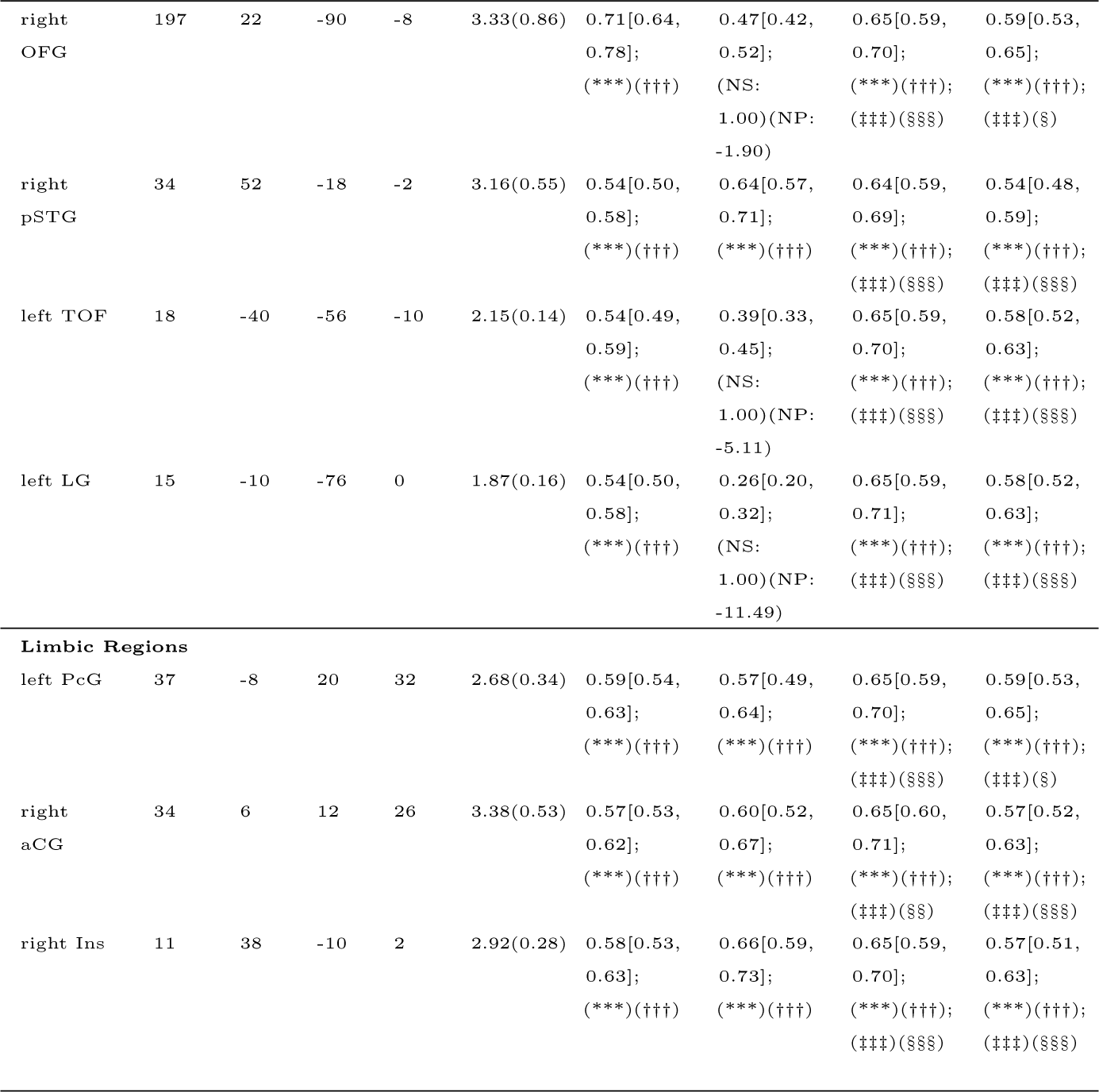
Single-cluster and ‘Virtual Lesion’ analysis of SA.

The single-cluster analysis enabled us to assess each cluster’s predictability in both intra- and inter-task scenarios, and to ascertain whether a single cluster alone was sufficient for prediction. As shown in Table 1 and Table 2, apart from 9 clusters in the SAS that could not predict conditions in the intra-task scenario, most clusters in the frontoparietal regions were sufficient for predicting outcomes in both intra-task and inter-task scenarios, and thus they exhibited similar representational patterns between FA and SA. Conversely, all clusters in the temporal/occipital regions displayed distinct representational patterns between FA and SA, because they showed predictive capabilities exclusively within the intra-task scenario. The single cluster analysis revealed the generalbility of the frontalparietal regions and the specificity of the temporal/occipital regions.

In addition, the ‘virtual lesion’ analysis allowed us to investigate the necessity of specific brain clusters by temporarily removing the cluster from the brain signatures. If the predictability of the remaining areas within the signature dropped to chance level after removing the cluster, it indicated that the cluster was necessary for making predictions. As shown in Table 1 and Table 2, for both FAS and SAS, the intra-task classification accuracy remained significantly higher than chance after removing each individual cluster. While the inter-task prediction was also unimpacted for FAS, the removal of some clusters in SAS could lead to chance-level prediction performance, though the significance was marginal. Additionally, when comparing the prediction accuracy after removing each single cluster with that of the entire brain signature, no clusters showed significant decrease in accuracy. These findings supported that no single cluster was necessary for predicting FA or SA.

To account for the potential confounding factor of varying voxel numbers among clusters, we conducted stepwise single-cluster analysis by randomly selecting the same significant voxel number for each cluster (Fig 4a). Regarding FA, the whole-brain signature consistently outperformed all individual clusters in both intra- and inter-task scenarios (for the complete trajectory of the whole-brain signature, see Supplementary Figure 3). As for SA, within the intra-task scenario, the most accurate prediction models in the single-cluster analysis were found in occipital regions, including right OcP, right OFG and right iLoC. The superior prediction performance compared to whole-brain single-cluster analysis might be attributed to the sparseness of significant voxels in the SAS, as the whole-brain SAS exhibited a prediction advantage when voxel numbers increased or in the ‘virtual lesion’ analysis.

**Fig 4.**
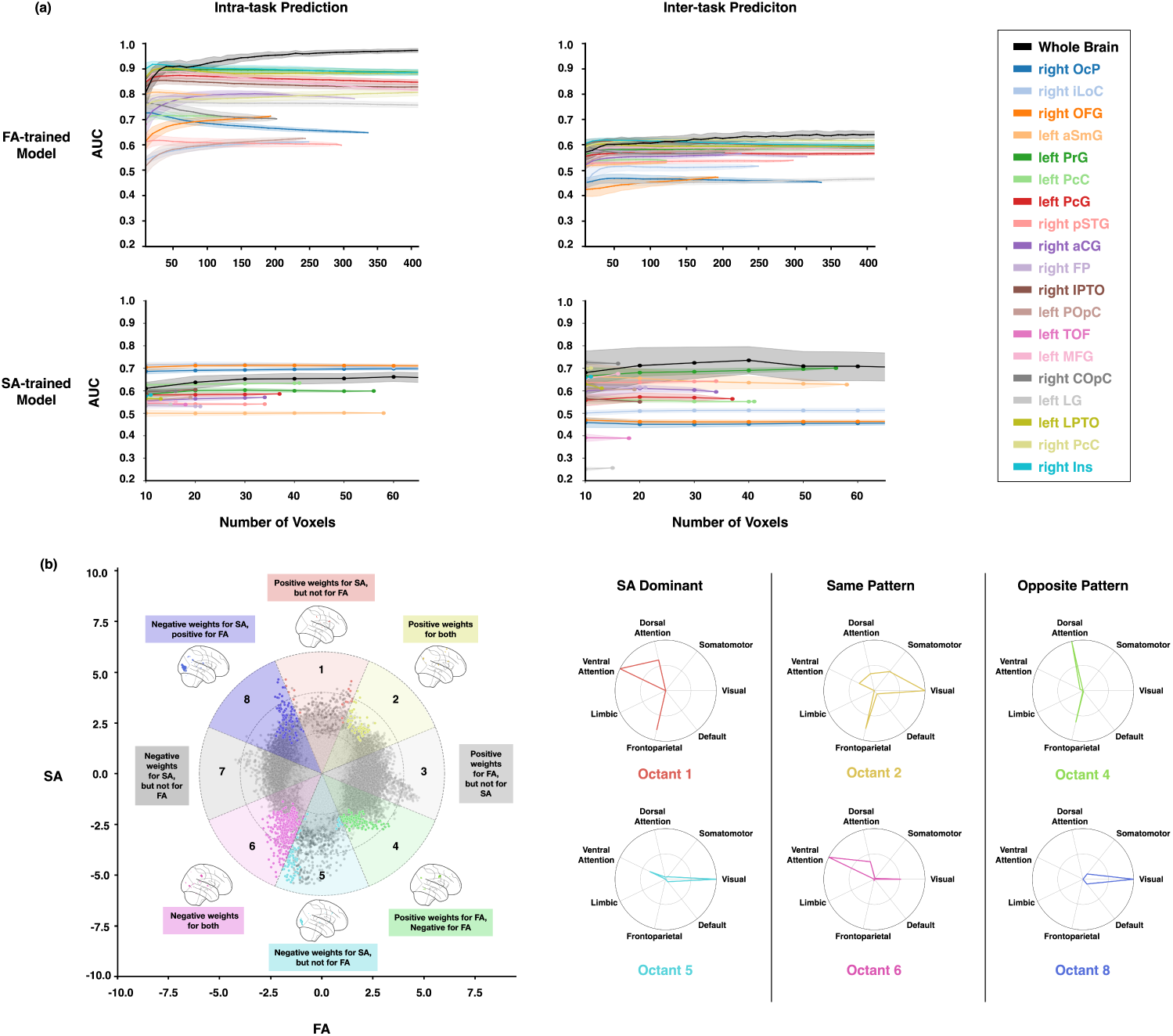
Single-cluster analysis and Voxel-level analysis. (a) illustrates the outcome of stepwise single-cluster analysis, where each cluster was tested on both intra-task and inter-task datasets in steps of 10 voxels. For each step, we randomly sampled voxels within the cluster 1,000 times to retrain classifiers and measured model performance using the AUC score. Shaded regions indicate the standard deviation. (b) depicts the whole-brain neural signatures of FA and SA on a 2-D plane with normalized absolute voxel values within each signature. The plane is divided into octants, categorizing voxel patterns into FA dominant (Octants 3 and 7), SA dominant (Octants 1 and 5), Same Pattern (Octants 2 and 6), and Opposite Pattern (Octants 4 and 8). Light gray dots represent voxels uniquely predictive for FA, while dark gray dots represent those uniquely predictive for SA. Colored dots indicate voxels predictive for both FA and SA. Glass brains display the brain regions of conjunction voxels. Radar plots show the normalized cortical coverage of significant voxels across cortical networks in each octant.

In short, the single-cluster analysis and the “virtual lesion” analysis revealed both shared and distinct representation patterns in FA and SA, along with the engagement of a distributed brain network in processing both forms of attention. While most individual conjunctive clusters could partially represent both FA and SA, none of them individually matched the effectiveness of the whole-brain signature in handling attention processing. Therefore, all the clusters were sufficient but not necessary to differentiate the two conditions in both forms of attention.

To further characterize the distributed nature of FAS and SAS, we performed a voxel-level analysis, plotting the voxels of both signatures together on a normalized 2-D plane (Fig 4b). The plane was evenly divided into octants, with voxels in each octants having the same attentional selectivity. Conjunction voxels were color-coded, FA-specific voxels were represented in light gray, and SA-specific voxels were shown in dark gray. Next, we conducted a comparative analysis of the cortical extent covered by conjunction voxels within cortical networks across the octants. The results, presented in radar plots (Fig 4b), revealed an intriguing discovery: each octant exhibited coverage of multiple networks, once again supporting the idea of a distributed representation of attention.

## Discussion

In this study, we utilized a fine-grained between-subject whole-brain multivariate machine-learning approach to explore the neural mechanisms underlying FA and SA. Our findings revealed robust brain signatures for both attentional processes respectively, which showed predictive capabilities not only within their respective tasks but also across tasks. This mutual predictivity suggested shared components between the FAS and SAS, while reduced inter-task prediction performance indicated the presence of distinct components within each signature. Moreover, we observed overlapping regions, consistent with previous univariate analyses, along with task-specific regions within the two brain signatures. Through single-cluster analysis and “virtual lesion” analysis on the overlapping clusters, we further revealed both shared and distinct representation patterns in FA and SA. This was evidenced by inter-task predictability in certain conjunctive clusters and its absence in others. Moreover, within the intra-task scenario, although each single cluster was sufficient for processing SA and FA, none of the clusters were necessary for either form of attention, supporting the distributed nature of these two brain signatures.

Unlike traditional brain mapping methods that focus on isolated core regions’ activation strengths or representational patterns, such as univariate activation analyses and single-subject level multivariate machine-learning approaches (e.g., MVPA, representational similarity pattern analyses), our study employed between-subject whole-brain multivariate machine-learning approaches to construct predictive models that maps whole-brain activity patterns to specific mental states or behaviors (in this study: FA conditions, or SA conditions). Despite having higher complexity and lower interpretability, these approaches integrate neural information across multiple distributed brain networks, resulting in sensitive brain signatures with greater predictive power for mind/behavior states (Chang et al., 2015; Kragel et al., 2018; Krishnan et al., 2016; Wager et al., 2013; F. Zhou et al., 2021). To mitigate overfitting risk and enabling the use of between-subject whole-brain multivariate machine-learning approaches, we included a substantial sample of 235 participants for fMRI scanning during FA and SA tasks, and validated the training models on a separate hold-out test dataset. We developed sensitive and reliable brain signatures of FA and SA respectively using the SVC algorithm, and validated them using the PCA-Logistic algorithm. The findings further offered a comprehensive understanding of the neural representation of FA and SA and their relation.

Previous fMRI studies using univariate activation analyses have identified overlapping activations in fronto-parietal-temporal attention-related reegions for both FA and SA (Galashan & Siemann, 2017; Giesbrecht et al., 2003; Wojciulik & Kanwisher, 1999), along with common attention-dependent activity modulation in visual areas (van Es et al., 2018). However, these findings relied on Boolean maps with p-value thresholds, considering conjunction areas as shared processing components. In addition, although single-subject level multivariate machine-learning approaches could identify searchlight or region of interests (ROIs) with different activity patterns across conditions, it lacked voxel-specific preferences. Our high spatial resolution brain signature, incorporating voxel-specific preferences, revealed common regions in dorsal and ventral attention networks and visual areas related to both attentional processes, including IPTO and SmG, consistently with previous research. Notably, uncorrelated weight patterns between FA and SA common clusters and reduced inter-task prediction accuracies, both indicate partial overlapping yet distinct neural representations, aligning with electrophysiological research (Bichot et al., 2015; Cohen & Maunsell, 2011; David et al., 2008; Goddard et al., 2022; Hayden & Gallant, 2005; McAdams & Maunsell, 2000; Ni & Maunsell, 2019). Our study offered a comprehensive whole-brain perspective on FA and SA, surpassing local neural population or neuron-level analysis.

An additional advancement of multivariate machine-learning approaches is the development of quantitative statistical models, providing advantages over qualitative models in univariate analyses. In detail, we assessed intra- and inter-task prediction performance through AUC scores at three levels: whole-brain, network, and cluster, enabling a robust evaluation of the similarity and distinctiveness of the two brain signatures.

At the whole-brain level, the FAS and SAS exhibited mutual predictability, albeit with reduced capacity in the inter-task scenario, suggesting the existence of both shared and distinct components. Furthermore, we evaluated network-level prediction performance using individual established cortical networks (Yeo et al., 2011). All seven networks demonstrated above-chance performance in the intra-task scenario, indicating the involvement of multiple distributed networks in both FA and SA. Specifically, the frontoparietal network exhibited the highest predictive performance for FA, while the visual network excelled in predicting SA, highlighting their respective prominence in the two attention processes.

At the finer cluster level, we employed single-cluster and ‘virtual lesion’ techniques to investigate the sufficiency and necessity of individual clusters for FA and SA. The single-cluster analysis revealed that the frontalparietal clusters were sufficient in both intra- and inter-task scenarios, while temporal/occipital clusters exhibited task-specificity. This disassociation of sufficiency implied similar top-down modulation from the frontoparietal regions, but with different modulation effects on sensory areas (Cohen & Maunsell, 2011; David et al., 2008; Goddard et al., 2022; Hayden & Gallant, 2005; McAdams & Maunsell, 2000; Ni & Maunsell, 2019; Patzwahl & Treue, 2009).

Additionaly, the ‘virtual lesion’ analysis enabled us to iteratively remove individual clusters from the whole-brain prediction model and evaluate their irreplaceable contributions. Surprisingly, no cluster was found to be necessary for FA or SA, as lesioning any cluster did not result in chance-level performance. These findings suggested a distributed brain functioning in both FA and SA. It was further supported by the voxel-level observation showing that the voxels with similar attentional selectivity spanned multiple networks.

The perspective ability of distributed representation of FA and SA, as revealed by machine-learning models with high predictability yet lower interpretability, challenges the traditional network-centric attention models that emphasize a few core regions, such as IPS, FEF, temporo-parietal junction (TPJ), and inferior frontal junction (IFJ), as primary centers for selective attention processing (Corbetta & Shulman, 2002, 2011; Petersen & Posner, 2012). This idea aligns with the emerging connectome-based predictive models, which have demonstrated effective individualized predictions of attentional and other mental/behavioral states through broadly distributed across-network connections (Finn et al., 2015; Gao et al., 2020; Rosenberg et al., 2016; Wu et al., 2020). Both perspectives emphasize the significance of large-scale, multivariate neural networks in contributing to cognition and behavior, supporting the notion that attentional processes may rely on distributed and interconnected brain networks (Bassett & Sporns, 2017; Mišíc & Sporns, 2016; Steinmetz et al., 2019; Urai et al., 2022). The shift from classic univariate (core region(s) corresponding to one specific behavior) to large-scale, multivariate neural network frameworks allows for advancements in linking brain elements to different cognition and behaviors (Bassett & Sporns, 2017). Further, it facilitates the development of more efficient neural modulation techniques compared to traditional modulation techniques focused on single core regions (Ching et al., 2012; Gu et al., 2015). These developments hold promise for enhancing or restoring attention abilities in individuals with attention-related disorders, such as autism spectrum disorder (ASD) and Attention-Deficit/Hyperactivity Disorder (ADHD).

In addition, by leveraging publicly available brain-wide gene expression atlases, such as the Allen Human Brain Atlas (AHBA) (Hawrylycz et al., 2012; Shen et al., 2012), the FAS and SAS can serve as valuable neuroimaging phenotypes for identifying attention-related gene expression. This approach opens up new possibilities to investigate the interplay between genetic expression profiles and the distributed FAS and SAS, exploring their associations with attentional states and potentially developing neurogenetic signatures of FA and SA (Bueichekú et al., 2020; Burt et al., 2018).

In summary, we successfully identified robust whole-brain neural representations of FA and SA (i.e., FAS and SAS). These two brain signatures exhibited both partial overlap and distinct representational characteristics. Moreover, they both demonstrated distributed representations across large-scale brain networks: all frontal/parietal clusters of the signatures were sufficient for both intra- and inter-task scenarios, while most temporal/occipital clusters showed task-specificity; none were deemed necessary for either FA or SA. These comprehensive understanding and direct comparison of the neural correlates in FA and SA, shed light on the coexistence of both shared and distinct mechanisms.

## Acknowledgments

This work was supported by the STI2030-Major Projects (2021ZD0203803), National Natural Science Foundation of China (32200840), National Key R&D Program of China (2019YFA0709503), China Postdoctoral Science Foundation (2022T150061, 2022M710435), and Fundamental Research Funds for the Central Universities.

