## Supplementary Information for "Shared and distinct neural signatures of feature and spatial attention"

### Supplementary Fig. 1

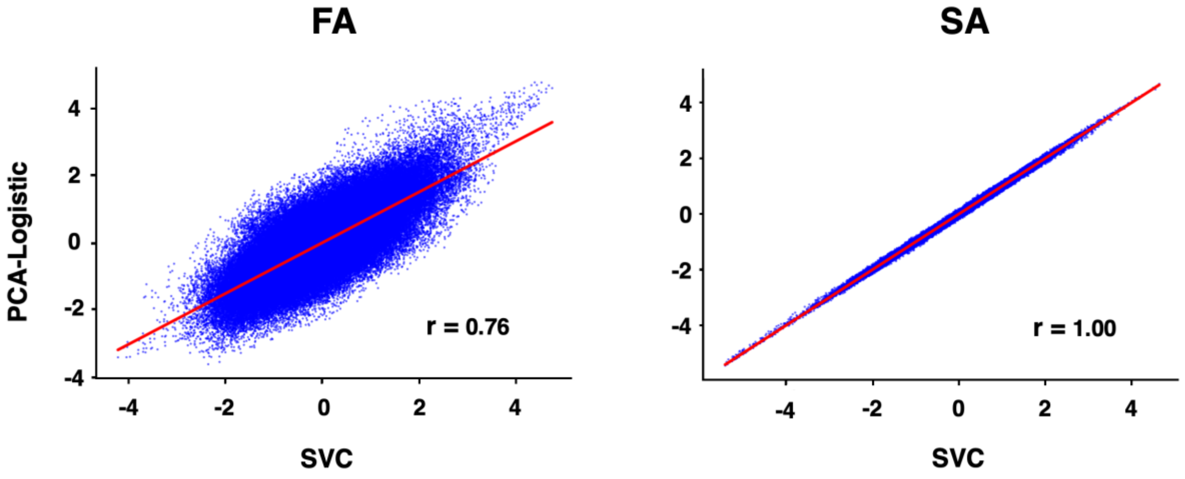


**Supplementary Fig. 1 Method Comparison between PCA-Logistic and SVC.** To assess the robustness of the SVC method employed in this study, we also utilized PCA-Logistic as a reference approach. We normalized the weights obtained from each method and represented them on a 2-D plane. Subsequently, we calculated the Pearson correlation for each pair of signatures. Notably, both cases demonstrated a significant correlation ($p < 0.001$), supporting the consistency and reliability of our findings.

**Supplementary Fig. 2**

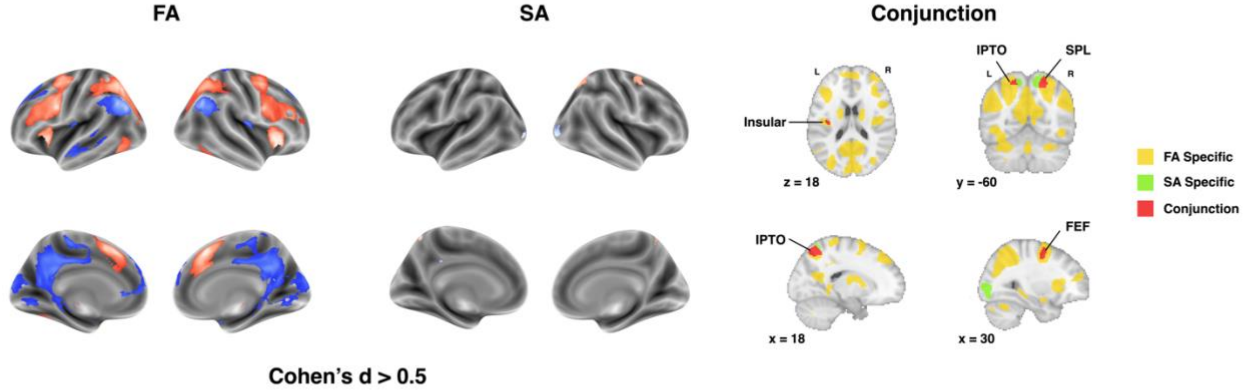


**Supplementary Fig. 2 Univariate Analysis.** Given the robustness of our study with a substantial sample size, effect size was utilized as the criterion for identifying significant voxels. We conducted paired-sample two-tailed t-tests on both FA and SA beta maps, and depicted voxels with an absolute Cohen's d value exceeding 0.5 (indicating a moderate effect size) in the figure. The resulting significant voxels were further classified into three categories: specific to FA, specific to SA, or showing a conjunction activation of FA and SA. IPTO: intraparietal/transverse occipital sulci, SPL: Superior Parietal Lobe, FEF: Frontal Eye Field.

**Supplementary Fig. 3**

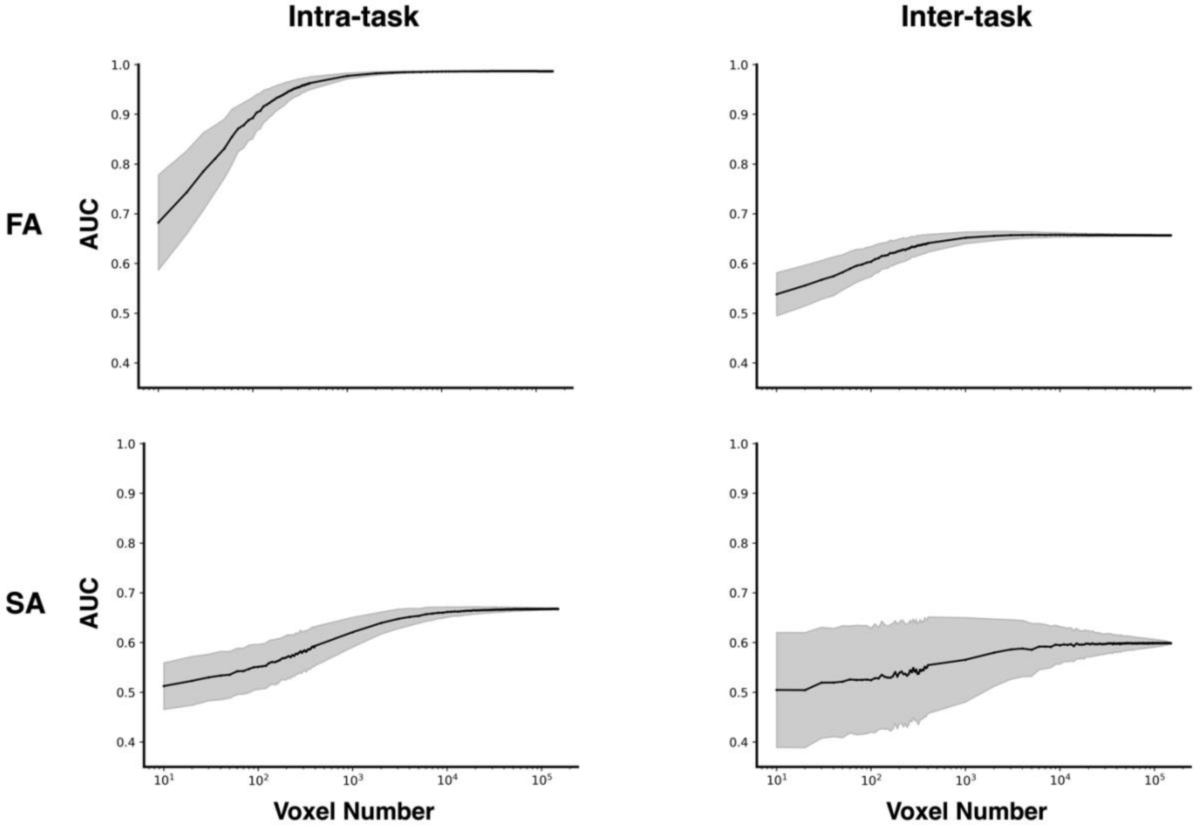


**Supplementary Fig. 3 Trajectory of Whole-Brain Single Cluster Analysis.** Voxels within the whole-brain signatures of FA and SA were analyzed at the step of 10 voxels. In each step, voxels were randomly sampled over 1,000 times, and prediction models were evaluated on both intra- and inter-tasks using AUC scores. The shaded regions represent the standard deviation.

### Supplementary Table 1

| | Difficult Tasks | Easy Tasks | t-value | effect size |
|---|---|---|---|---|
| | | **Feature-based Attention** | | |
| | Feature Task(SE) | Conjunction Task(SE) | | |
| d' | 4.27±0.02 | 2.09±0.04 | 55.18(***) | 4.92 |
| RT | 451.57±2.97 | 584.42±3.38 | -36.29(***) | 2.75 |
| | | **Space-based Attention** | | |
| | Central Task(SE) | Peripheral Task(SE) | | |
| d' | 3.82±0.04 | 3.07±0.03 | 16.74(***) | 1.33 |
| RT | 490.93±2.53 | 497.54±2.67 | -2.78(**) | 0.17 |

**Supplementary Table 1 Behavioral Results of FA and SA experiments.** The d' index quantifies participants' ability to discriminate signals from distractors, calculated as $d' = Z(hit\ rate) - Z(false\ alarm\ rate)$. Reaction time (RT) measures the elapsed time from signal presentation to key press. Asterisks denote p-value: ***, $p < 0.001$; **, $p < 0.01$.

### Supplementary Table 2

| | Test Dataset: FA | | | | | Test Dataset: SA | | | | |
|---|---|---|---|---|---|---|---|---|---|---|
| Neural Signatures | accuracy | precision | recall | f-measure | AUC | accuracy | precision | recall | f-measure | AUC |
| FA | 0.95[0.82, 0.98] | 0.96[0.92, 1.00] | 0.94[0.88, 0.99] | 0.95[0.91, 0.98] | 0.99[0.97, 1.00] | 0.62[0.54, 0.69] | 0.61[0.54, 0.69] | 0.63[0.52, 0.74] | 0.62[0.54, 0.70] | 0.66[0.58, 0.73] |
| SA | 0.56[0.50, 0.63] | 0.57[0.50, 0.64] | 0.54[0.43, 0.65] | 0.55[0.47, 0.63] | 0.60[0.54, 0.66] | 0.62[0.56, 0.67] | 0.62[0.56, 0.68] | 0.61[0.51, 0.72] | 0.61[0.55, 0.68] | 0.67[0.62, 0.72] |

**Supplementary Table 2 Model Performance.** The model performances of both neural signatures were evaluated using a hold-out test dataset with multiple metrics, including accuracy, precision, recall, f-measure and AUC. The table presents both the mean values and 95% CIs, which were constructed through 10,000 bootstrap resampling iterations.

### Supplementary Table 3

| Cluster | Voxel Number | Pearson's r | p-value |
|---|---|---|---|
| right iLOC | 107 | 0.15 | 0.13 |
| right OcP | 101 | 0.24 | 0.02 |
| right OFG | 71 | -0.49 | < 0.001 |
| left PrG | 54 | -0.39 | < 0.001 |
| left aSMG | 39 | -0.15 | 0.38 |
| right IPTO | 20 | 0.23 | 0.33 |
| left LG | 15 | -0.33 | 0.23 |
| right aCG | 13 | 0.11 | 0.73 |
| left IPTO | 13 | -0.11 | 0.73 |
| right PcC | 10 | -0.01 | 0.97 |
| right Ins | 10 | -0.32 | 0.37 |
| right COpC | 8 | -0.02 | 0.97 |
| right pSTG | 7 | -0.38 | 0.4 |
| left POpC | 6 | 0.04 | 0.93 |
| left PcG | 5 | 0.84 | 0.07 |

**Supplementary Table 3 Correlation of each conjunction region's representational patterns between FA and SA.** Complete names of the brain regions are referenced in Supplementary Table 5.

### Supplementary Table 4

| Clusters | Number of Voxels | MNI Peak Coordinates | | | z-score | Single-Cluster | | Lesion | |
|---|---|---|---|---|---|---|---|---|---|
| | | x | y | z | Mean(SD) | Intra Task | Inter Task | Intra Task | Inter Task |
| left IPTO | 3343 | -20 | -74 | 48 | 1.86(0.77) | 0.89[0.84, 0.93]; (***) | 0.59[0.53, 0.65]; (*) | 0.99[0.98, 1.00]; (***); (NS: 1.00) | 0.66[0.59, 0.73]; (***); (NS: 1.00) |
| right IPTO | 1711 | 26 | -58 | 46 | 1.64(0.50) | 0.83[0.78, 0.88]; (***) | 0.6[0.54, 0.65]; (*) | 0.99[0.98, 1.00]; (***); (NS: 1.00) | 0.66[0.59, 0.73]; (***); (NS: 1.00) |
| left PrG | 1646 | -36 | -6 | 50 | 1.61(0.54) | 0.88[0.84, 0.93]; (***) | 0.59[0.53, 0.64]; (*) | 0.99[0.98, 1.00]; (***); (NS: 1.00) | 0.67[0.59, 0.74]; (***); (NS: 1.00) |
| right PcG | 1330 | 6 | 22 | 42 | 1.56(0.51) | 0.86[0.81, 0.90]; (***) | 0.61[0.55, 0.67]; (**) | 0.99[0.97, 1.00]; (***); (NS: 1.00) | 0.67[0.59, 0.74]; (***); (NS: 1.00) |
| left Ins | 1233 | -32 | 22 | 0 | 1.87(0.46) | 0.9[0.85, 0.94]; (***) | 0.61[0.55, 0.67]; (**) | 0.99[0.98, 1.00]; (***); (NS: 1.00) | 0.66[0.59, 0.74]; (***); (NS: 1.00) |
| left PoG | 1186 | -44 | -32 | 62 | 1.59(0.44) | 0.8[0.74, 0.85]; (***) | 0.57[0.52, 0.61]; (NS: 0.07) | 0.99[0.98, 1.00]; (***); (NS: 1.00) | 0.66[0.58, 0.73]; (***); (NS: 1.00) |
| right Ins | 1183 | 36 | -20 | 0 | 1.60(0.47) | 0.89[0.85, 0.93]; (***) | 0.61[0.55, 0.67]; (*) | 0.99[0.98, 1.00]; (***); (NS: 1.00) | 0.66[0.59, 0.74]; (***); (NS: 1.00) |
| right AG | 1153 | 62 | -48 | 42 | 1.47(0.41) | 0.81[0.75, 0.87]; (***) | 0.56[0.52, 0.60]; (NS: 0.10) | 0.99[0.98, 1.00]; (***); (NS: 1.00) | 0.67[0.60, 0.74]; (***); (NS: 1.00) |
| left SFG | 1055 | -2 | 26 | 46 | 1.66(0.46) | 0.91[0.87, 0.95]; (***) | 0.56[0.50, 0.62]; (NS: 0.10) | 0.99[0.98, 1.00]; (***); (NS: 1.00) | 0.67[0.59, 0.74]; (***); (NS: 1.00) |
| right PrG | 1013 | 36 | -6 | 66 | 1.27(0.38) | 0.85[0.81, 0.90]; (***) | 0.63[0.57, 0.69]; (**) | 0.99[0.98, 1.00]; (***); (NS: 1.00) | 0.67[0.59, 0.74]; (***); (NS: 1.00) |
| right PcC | 874 | 12 | -60 | 46 | 1.50(0.38) | 0.82[0.77, 0.87]; (***) | 0.62[0.57, 0.67]; (**) | 0.99[0.98, 1.00]; (***); (NS: 1.00) | 0.67[0.60, 0.74]; (***); (NS: 1.00) |
| left LG | 807 | -26 | -54 | -8 | 1.60(0.41) | 0.75[0.70, 0.80]; (***) | 0.48[0.44, 0.52]; (NS: 0.66) | 0.99[0.98, 1.00]; (***); (NS: 1.00) | 0.67[0.59, 0.74]; (***); (NS: 1.00) |
| right CGp | 785 | 8 | -44 | 6 | 1.37(0.41) | 0.74[0.69, 0.80]; (***) | 0.57[0.52, 0.62]; (NS: 0.07) | 0.99[0.98, 1.00]; (***); (NS: 1.00) | 0.66[0.59, 0.74]; (***); (NS: 1.00) |
| right PoG | 769 | 48 | -20 | 60 | 1.56(0.44) | 0.73[0.66, 0.80]; (***) | 0.58[0.53, 0.63]; (*) | 0.99[0.98, 1.00]; (***); (NS: 1.00) | 0.67[0.59, 0.74]; (***); (NS: 1.00) |
| right MFG | 760 | 38 | 32 | 18 | 1.68(0.54) | 0.81[0.76, 0.87]; (***) | 0.55[0.51, 0.59]; (NS: 0.14) | 0.99[0.98, 1.00]; (***); (NS: 1.00) | 0.67[0.60, 0.74]; (***); (NS: 1.00) |
| left CGp | 747 | -12 | -38 | 38 | 1.42(0.47) | 0.76[0.70, 0.81]; (***) | 0.57[0.52, 0.63]; (NS: 0.06) | 0.99[0.98, 1.00]; (***); (NS: 1.00) | 0.67[0.59, 0.74]; (***); (NS: 1.00) |
| left SPL | 740 | -26 | -58 | 46 | 1.65(0.50) | 0.82[0.77, 0.87]; (***) | 0.57[0.50, 0.63]; (NS: 0.07) | 0.99[0.98, 1.00]; (***); (NS: 1.00) | 0.66[0.59, 0.74]; (***); (NS: 1.00) |
| left MFG | 689 | -42 | 28 | 30 | 1.33(0.40) | 0.8[0.74, 0.86]; (***) | 0.58[0.52, 0.64]; (*) | 0.99[0.98, 1.00]; (***); (NS: 1.00) | 0.67[0.59, 0.74]; (***); (NS: 1.00) |
| left SmGp | 677 | -54 | -50 | 48 | 1.69(0.36) | 0.83[0.77, 0.89]; (***) | 0.55[0.51, 0.59]; (NS: 0.14) | 0.99[0.98, 1.00]; (***); (NS: 1.00) | 0.67[0.59, 0.74]; (***); (NS: 1.00) |

| | | | | | | | | | |
|---|---|---|---|---|---|---|---|---|---|
| left PcG | 664 | -2 | 14 | 52 | 1.75(0.48) | 0.84[0.79, 0.89]; (***) | 0.57[0.52, 0.62]; (NS: 0.07) | 0.99[0.98, 1.00]; (***); (NS: 1.00) | 0.67[0.59, 0.74]; (***); (NS: 1.00) |
| right MTGtp | 614 | 62 | -52 | 6 | 2.00(0.61) | 0.63[0.57, 0.69]; (**) | 0.56[0.51, 0.60]; (NS: 0.10) | 0.99[0.98, 1.00]; (***); (NS: 1.00) | 0.66[0.59, 0.74]; (***); (NS: 1.00) |
| right SPL | 415 | 40 | -38 | 56 | 1.28(0.31) | 0.81[0.75, 0.87]; (***) | 0.6[0.54, 0.65]; (*) | 0.99[0.98, 1.00]; (***); (NS: 1.00) | 0.66[0.59, 0.74]; (***); (NS: 1.00) |
| right SMC | 392 | 6 | -10 | 62 | 1.36(0.37) | 0.81[0.76, 0.87]; (***) | 0.59[0.54, 0.65]; (*) | 0.99[0.98, 1.00]; (***); (NS: 1.00) | 0.67[0.60, 0.74]; (***); (NS: 1.00) |
| left IFGpo | 369 | -44 | 14 | 18 | 1.48(0.50) | 0.83[0.78, 0.88]; (***) | 0.55[0.51, 0.60]; (NS: 0.14) | 0.99[0.98, 1.00]; (***); (NS: 1.00) | 0.67[0.59, 0.74]; (***); (NS: 1.00) |
| left OcP | 368 | -6 | -94 | -6 | 1.42(0.36) | 0.76[0.70, 0.81]; (***) | 0.49[0.46, 0.53]; (NS: 0.58) | 0.99[0.98, 1.00]; (***); (NS: 1.00) | 0.67[0.59, 0.74]; (***); (NS: 1.00) |
| left aCG | 350 | -6 | 4 | 36 | 1.55(0.43) | 0.82[0.76, 0.88]; (***) | 0.58[0.54, 0.63]; (*) | 0.99[0.98, 1.00]; (***); (NS: 1.00) | 0.66[0.59, 0.73]; (***); (NS: 1.00) |
| left OFG | 342 | -22 | -76 | -6 | 1.64(0.37) | 0.77[0.71, 0.82]; (***) | 0.53[0.49, 0.57]; (NS: 0.26) | 0.99[0.98, 1.00]; (***); (NS: 1.00) | 0.67[0.59, 0.74]; (***); (NS: 1.00) |
| right aSmG | 336 | 60 | -34 | 40 | 1.81(0.49) | 0.65[0.60, 0.70]; (***) | 0.51[0.47, 0.55]; (NS: 0.41) | 0.99[0.98, 1.00]; (***); (NS: 1.00) | 0.67[0.59, 0.74]; (***); (NS: 1.00) |
| right OcP | 336 | 28 | -94 | 12 | 1.52(0.35) | 0.65[0.62, 0.69]; (***) | 0.45[0.41, 0.49]; (NS: 0.86) | 0.99[0.98, 1.00]; (***); (NS: 1.00) | 0.67[0.59, 0.74]; (***); (NS: 1.00) |
| right FP | 316 | 28 | 40 | 42 | 1.28(0.31) | 0.78[0.72, 0.85]; (***) | 0.55[0.51, 0.60]; (NS: 0.14) | 0.99[0.98, 1.00]; (***); (NS: 1.00) | 0.67[0.59, 0.74]; (***); (NS: 1.00) |
| right pSTG | 297 | 50 | -30 | 4 | 1.45(0.23) | 0.6[0.53, 0.67]; (*) | 0.54[0.50, 0.58]; (NS: 0.20) | 0.99[0.98, 1.00]; (***); (NS: 1.00) | 0.67[0.59, 0.74]; (***); (NS: 1.00) |
| right LG | 263 | 16 | -78 | -10 | 1.27(0.26) | 0.75[0.70, 0.81]; (***) | 0.53[0.48, 0.57]; (NS: 0.26) | 0.99[0.98, 1.00]; (***); (NS: 1.00) | 0.67[0.59, 0.74]; (***); (NS: 1.00) |
| right iLOC | 249 | 32 | -84 | -2 | 1.92(0.33) | 0.61[0.57, 0.65]; (**) | 0.52[0.47, 0.56]; (NS: 0.34) | 0.99[0.98, 1.00]; (***); (NS: 1.00) | 0.66[0.59, 0.73]; (***); (NS: 1.00) |
| right CC | 244 | 4 | -88 | 20 | 1.27(0.24) | 0.74[0.68, 0.79]; (***) | 0.51[0.48, 0.54]; (NS: 0.41) | 0.99[0.98, 1.00]; (***); (NS: 1.00) | 0.67[0.59, 0.74]; (***); (NS: 1.00) |
| left POpC | 244 | -46 | -28 | 16 | 1.74(0.42) | 0.63[0.56, 0.69]; (**) | 0.58[0.54, 0.63]; (*) | 0.99[0.98, 1.00]; (***); (NS: 1.00) | 0.67[0.59, 0.74]; (***); (NS: 1.00) |
| left pSTG | 241 | -60 | -30 | 6 | 1.46(0.32) | 0.63[0.56, 0.69]; (**) | 0.52[0.49, 0.56]; (NS: 0.33) | 0.99[0.98, 1.00]; (***); (NS: 1.00) | 0.67[0.59, 0.74]; (***); (NS: 1.00) |
| left pPaG | 233 | -14 | -34 | -16 | 2.10(0.54) | 0.66[0.60, 0.73]; (***) | 0.58[0.54, 0.63]; (*) | 0.99[0.98, 1.00]; (***); (NS: 1.00) | 0.67[0.60, 0.74]; (***); (NS: 1.00) |
| right aCG | 225 | 0 | 14 | 22 | 1.73(0.50) | 0.8[0.74, 0.86]; (***) | 0.57[0.53, 0.61]; (NS: 0.06) | 0.99[0.98, 1.00]; (***); (NS: 1.00) | 0.67[0.59, 0.74]; (***); (NS: 1.00) |
| left IFGpt | 213 | -54 | 28 | 6 | 1.25(0.35) | 0.8[0.74, 0.85]; (***) | 0.52[0.48, 0.56]; (NS: 0.33) | 0.99[0.98, 1.00]; (***); (NS: 1.00) | 0.67[0.59, 0.74]; (***); (NS: 1.00) |
| right SFG | 205 | 26 | 2 | 56 | 1.51(0.48) | 0.81[0.74, 0.87]; (***) | 0.61[0.54, 0.67]; (**) | 0.99[0.98, 1.00]; (***); (NS: 1.00) | 0.67[0.60, 0.74]; (***); (NS: 1.00) |
| left COpC | 204 | -52 | -6 | 12 | 1.52(0.28) | 0.71[0.65, 0.77]; (***) | 0.53[0.49, 0.58]; (NS: 0.26) | 0.99[0.98, 1.00]; (***); (NS: 1.00) | 0.67[0.59, 0.74]; (***); (NS: 1.00) |
| right COpC | 202 | 40 | -14 | 22 | 1.47(0.39) | 0.7[0.64, 0.77]; (***) | 0.57[0.53, 0.61]; (NS: 0.07) | 0.99[0.98, 1.00]; (***); (NS: 1.00) | 0.67[0.59, 0.74]; (***); (NS: 1.00) |
| left FOpC | 195 | -42 | 10 | 8 | 1.71(0.42) | 0.8[0.74, 0.86]; (***) | 0.55[0.51, 0.59]; (NS: 0.14) | 0.99[0.98, 1.00]; (***); (NS: 1.00) | 0.67[0.59, 0.74]; (***); (NS: 1.00) |

| | | | | | | | | | |
|---|---|---|---|---|---|---|---|---|---|
| right OFG | 193 | 20 | -86 | -6 | 1.72(0.31) | 0.72[0.66, 0.77]; (***) | 0.47[0.41, 0.53]; (NS: 0.73) | 0.99[0.98, 1.00]; (***); (NS: 1.00) | 0.67[0.59, 0.74]; (***); (NS: 1.00) |
| right pSmG | 152 | 68 | -40 | 18 | 1.18(0.25) | 0.72[0.65, 0.79]; (***) | 0.5[0.46, 0.54]; (NS: 0.50) | 0.99[0.98, 1.00]; (***); (NS: 1.00) | 0.67[0.59, 0.74]; (***); (NS: 1.00) |
| left SMC | 148 | -2 | 4 | 58 | 1.08(0.24) | 0.83[0.78, 0.89]; (***) | 0.63[0.57, 0.69]; (**) | 0.99[0.98, 1.00]; (***); (NS: 1.00) | 0.67[0.59, 0.74]; (***); (NS: 1.00) |
| left pMTGt | 131 | -56 | -50 | 0 | 1.20(0.23) | 0.74[0.68, 0.80]; (***) | 0.56[0.51, 0.61]; (NS: 0.09) | 0.99[0.98, 1.00]; (***); (NS: 1.00) | 0.67[0.59, 0.74]; (***); (NS: 1.00) |
| right pPaG | 130 | 16 | -30 | -8 | 1.69(0.43) | 0.69[0.62, 0.76]; (***) | 0.54[0.49, 0.58]; (NS: 0.20) | 0.99[0.98, 1.00]; (***); (NS: 1.00) | 0.67[0.59, 0.74]; (***); (NS: 1.00) |
| left aSmG | 122 | -60 | -34 | 40 | 1.59(0.30) | 0.8[0.74, 0.85]; (***) | 0.53[0.49, 0.58]; (NS: 0.25) | 0.99[0.98, 1.00]; (***); (NS: 1.00) | 0.67[0.59, 0.74]; (***); (NS: 1.00) |
| left PcC | 122 | -8 | -62 | 44 | 1.39(0.20) | 0.71[0.65, 0.77]; (***) | 0.54[0.50, 0.58]; (NS: 0.20) | 0.99[0.98, 1.00]; (***); (NS: 1.00) | 0.67[0.60, 0.74]; (***); (NS: 1.00) |
| left pITGt | 104 | -42 | -54 | -10 | 1.41(0.27) | 0.72[0.67, 0.78]; (***) | 0.46[0.42, 0.50]; (NS: 0.80) | 0.99[0.98, 1.00]; (***); (NS: 1.00) | 0.67[0.60, 0.74]; (***); (NS: 1.00) |
| left pMTG | 100 | -60 | -30 | -10 | 1.09(0.20) | 0.61[0.55, 0.68]; (**) | 0.56[0.53, 0.59]; (NS: 0.10) | 0.99[0.98, 1.00]; (***); (NS: 1.00) | 0.67[0.59, 0.74]; (***); (NS: 1.00) |
| right IFGpo | 99 | 42 | 12 | 22 | 1.10(0.37) | 0.79[0.73, 0.84]; (***) | 0.54[0.50, 0.58]; (NS: 0.19) | 0.99[0.98, 1.00]; (***); (NS: 1.00) | 0.67[0.59, 0.74]; (***); (NS: 1.00) |
| right POpC | 94 | 38 | -32 | 22 | 1.55(0.23) | 0.72[0.65, 0.78]; (***) | 0.57[0.52, 0.61]; (NS: 0.07) | 0.99[0.98, 1.00]; (***); (NS: 1.00) | 0.67[0.59, 0.74]; (***); (NS: 1.00) |
| left AG | 91 | -62 | -54 | 22 | 1.37(0.27) | 0.78[0.72, 0.83]; (***) | 0.56[0.51, 0.60]; (NS: 0.10) | 0.99[0.98, 1.00]; (***); (NS: 1.00) | 0.67[0.59, 0.74]; (***); (NS: 1.00) |
| left iLOC | 82 | -50 | -74 | 2 | 1.44(0.33) | 0.7[0.66, 0.75]; (***) | 0.43[0.40, 0.47]; (NS: 0.94) | 0.99[0.98, 1.00]; (***); (NS: 1.00) | 0.67[0.59, 0.74]; (***); (NS: 1.00) |
| left FP | 78 | -20 | 56 | 14 | 1.22(0.23) | 0.75[0.69, 0.82]; (***) | 0.58[0.53, 0.64]; (*) | 0.99[0.98, 1.00]; (***); (NS: 1.00) | 0.67[0.59, 0.74]; (***); (NS: 1.00) |
| right TOF | 61 | 40 | -46 | -18 | 1.41(0.29) | 0.63[0.57, 0.69]; (**) | 0.47[0.43, 0.51]; (NS: 0.74) | 0.99[0.98, 1.00]; (***); (NS: 1.00) | 0.67[0.59, 0.74]; (***); (NS: 1.00) |
| right H1/H2 | 60 | 52 | -12 | 2 | 1.48(0.20) | 0.67[0.61, 0.73]; (***) | 0.55[0.50, 0.60]; (NS: 0.14) | 0.99[0.98, 1.00]; (***); (NS: 1.00) | 0.67[0.59, 0.74]; (***); (NS: 1.00) |
| right PT | 57 | 42 | -30 | 16 | 1.64(0.29) | 0.63[0.55, 0.71]; (**) | 0.53[0.49, 0.57]; (NS: 0.26) | 0.99[0.98, 1.00]; (***); (NS: 1.00) | 0.67[0.59, 0.74]; (***); (NS: 1.00) |
| right FOC | 54 | 36 | 18 | -18 | 1.67(0.25) | 0.79[0.73, 0.85]; (***) | 0.55[0.51, 0.60]; (NS: 0.14) | 0.99[0.98, 1.00]; (***); (NS: 1.00) | 0.67[0.59, 0.74]; (***); (NS: 1.00) |
| left aSTG | 27 | -48 | 0 | -22 | 1.61(0.35) | 0.69[0.63, 0.76]; (***) | 0.58[0.53, 0.63]; (*) | 0.99[0.98, 1.00]; (***); (NS: 1.00) | 0.67[0.60, 0.74]; (***); (NS: 1.00) |
| left FOC | 20 | -38 | 18 | -12 | 1.54(0.19) | 0.67[0.61, 0.74]; (***) | 0.56[0.51, 0.60]; (NS: 0.09) | 0.99[0.98, 1.00]; (***); (NS: 1.00) | 0.67[0.59, 0.74]; (***); (NS: 1.00) |
| left TOF | 11 | -40 | -58 | -14 | 1.35(0.14) | 0.68[0.62, 0.74]; (***) | 0.43[0.39, 0.48]; (NS: 0.93) | 0.99[0.98, 1.00]; (***); (NS: 1.00) | 0.67[0.59, 0.74]; (***); (NS: 1.00) |

**Supplementary Table 4 Complete table of single-cluster and 'virtual lesion' analysis in FA signature.** FA clusters were obtained through watershed algorithm with the minimum voxel size at 10. 'Intra-task' refers to testing the model on the same task it was trained for. ‘Inter-task’ refers to testing the model on a different task that it has not been trained on. Model performance was measured by AUC. We constructed 95% CI through bootstrap over 10,000 times. For single-cluster analysis, model performance was evaluated with permutation over 10, 000 times. For ‘virtual lesion’ analysis, the impact of the virtual lesion was assessed through both model performance and the reduction in model performance compared to the whole-brain condition. Full names of the brain regions should refer to Supplementary Table 5.

Asterisks indicate the p-value of statistical inference for model performance against chance level ($AUC = 0.5$). *, $p < 0.05$; **, $p < 0.01$; ***, $p < 0.001$; NS, not significant, $p > 0.05$, with the specific p-value indicated.

Daggers indicate the p-value of statistical inference for model performance drop ($\Delta AUC = 0$). †, $p < 0.05$; ††, $p < 0.01$; †††, $p < 0.001$; NS, not significant, $p > 0.05$, with the specific p-value indicated. Note in the table no cluster lesion causes significant performance drop.

### Supplementary Table 5

| Abbreviation | Region Name |
|---|---|
| FP | Frontal Pole |
| Ins | Insular Cortex |
| SFG | Superior Frontal Gyrus |
| MFG | Middle Frontal Gyrus |
| IFGpt | Inferior Frontal Gyrus, pars triangularis |
| IFGpo | Inferior Frontal Gyrus, pars opercularis |
| PrG | Precentral Gyrus |
| aSTG | Superior Temporal Gyrus, anterior division |
| pSTG | Superior Temporal Gyrus, posterior division |
| pMTG | Middle Temporal Gyrus, posterior division |
| tpMTG | Middle Temporal Gyrus, temporooccipital part |
| tpITG | Inferior Temporal Gyrus, temporooccipital part |
| PoG | Postcentral Gyrus |
| SPL | Superior Parietal Lobule |
| aSmG | Supramarginal Gyrus, anterior division |
| pSmG | Supramarginal Gyrus, posterior division |
| AG | Angular Gyrus |
| IPTO | Intraparietal/transverse occipital sulci |
| iLOC | Lateral Occipital Cortex, inferior division |
| SMC | Supplementary Motor Cortex |
| PcG | Paracingulate Gyrus |
| aCG | Cingulate Gyrus, anterior division |
| pCG | Cingulate Gyrus, posterior division |
| PcC | Precuneus Cortex |
| CC | Cuneal Cortex |
| FOC | Frontal Orbital Cortex |
| pPaG | Parahippocampal Gyrus, posterior division |
| LG | Lingual Gyrus |
| TOF | Temporal Occipital Fusiform Cortex |
| OFG | Occipital Fusiform Gyrus |
| FOpC | Frontal Operculum Cortex |
| COpC | Central Opercular Cortex |
| POpC | Parietal Operculum Cortex |
| H1/H2 | Heschl's Gyrus (includes H1 and H2) |
| PT | Planum Temporale |
| OcP | Occipital Pole |

**Supplementary Table 5 The complete names corresponding to the abbreviations of brain regions.**
